# Discovery and CryoEM Structure of FPM13, a Periplasmic Metalloprotein Unique to *Francisella*

**DOI:** 10.1101/2025.05.08.652791

**Authors:** Daniel L. Clemens, Bai-Yu Lee, Xiaoyu Liu, Z. Hong Zhou, Marcus A. Horwitz

## Abstract

We report the cryoEM structure of the *Francisella* protein FTN_1118, identifying it as a novel 13 kDa periplasmic protein unique to the *Francisella* genus, which we now designate FPM13 (*Francisella* Periplasmic Metalloprotein, 13 kDa) based on its structural and biochemical properties. FPM13 was serendipitously identified during purification of *Francisella* type VI secretion system (T6SS) effector proteins, co-purifying with them. Its identity, initially unknown, was established using the novel cryoID method. The structure reveals a symmetrical, donut-shaped 18-mer with 9-fold dihedral symmetry, formed by two stacked nonamers head-to-head. It measures ∼8 nm both in height and in outer diameter, and has a 3.5 nm central channel. The complex features a double-layered wall with an inner β-sheet core and an outer α-helical shell. Each monomer adopts a compact fold comprising an N-terminus β-strand, an α-helix and two additional β strands at the C-terminus. The assembly is stabilized by inter-ring loop interactions and hydrophobic and electrostatic contacts between neighboring subunits. Biochemical analyses, as shown by APEX-biotinylation and Triton X-114 phase partitioning, confirmed that FPM13 is a soluble periplasmic protein. ICP-MS demonstrated that FPM13 binds iron and copper. Deletion of FPM13 in *Francisella novicida* strains caused no growth defects in macrophages or mice but show increased copper sensitivity under iron-depleted conditions, suggesting a role for FPM13 in metal transport or detoxification.

## INTRODUCTION

Tularemia, also known as “rabbit fever”, is a potentially fatal bacterial zoonotic disease whose etiologic agent is *Francisella tularensis,* a gram negative coccobacillus and member of the *Francisellaceae* family. The *Francisellaceae* species are widely distributed in nature and cause natural infections in a wide range of animals, including mammals, birds, amphibians, fish, mollusks, insects, and protists. Some species are obligate pathogens of animals and humans (e.g., *F. tularensis*), whereas other species are present in the environment and are opportunistic pathogens of humans (*F. novicida* and *F. philomiragia*). Other species of *Francisellaceae* have been associated only with animals (e.g., *F. noatunensis*) or protists (e.g., *F. endociliophora*), and others (e.g., *F. persica*) are endosymbionts that lives only in ticks.

In nature, *F. tularensis* infects lagomorphs or voles, and blood sucking insects (e.g., ticks, deer flies, and mosquitoes) serve as intermediate hosts that can transmit the infection to humans. *F. tularensis* is highly infectious; inoculation^1^ or inhalation^2^ of as few as 10 or 25 organisms, respectively, can cause potentially lethal infection in humans. Transmission of tularemia can occur from the bite of an infected tick, deerfly or other insects; from handling infected animal carcasses; from eating or drinking contaminated food or water; or from inhaling the bacteria^2^.

Macrophages are the primary host cells for *F. tularensis* in animals^3^. After uptake by macrophages via the novel mechanism of looping phagocytosis^4^, *F. tularensis* initially resides in a phagosome that resists fusion with lysosomes and fails to acquire acid hydrolases, such as cathepsin D^5^. Within hours, the bacterium escapes from its phagosome and multiplies extensively in the macrophage cytosol^6^. The *Francisella* Pathogenicity Island (FPI), a cluster of ∼17 genes, is a major virulence determinant that encodes a Type 6 Secretion System (T6SS)^7^. Disruption of almost any of the FPI genes renders the bacterium unable to escape the phagosome, multiply in host macrophages, and cause disease^8^. Because of this, a major focus of our research has been to purify and determine the structure and function of the components of the *Francisella* T6SS, especially its secreted effector proteins. Because it shares the same intracellular lifestyle and is genetically very similar to the more pathogenic *F. tularensis*, many of our studies are conducted in the closely related species *F. novicida*, which was first isolated from saltwater taken from Great Salt Lake, Utah^9^. *F. novicida* rarely can cause infections in humans, including an outbreak in a Louisiana prison, with ice machines being identified as the common source^10^.

In this study, we serendipitously determined the cryoEM structure of a native *F. novicida* protein complex during the purification of T6SS effector proteins and subsequently identified using the cryoID approach as an 18-mer complex of FTN_1118, a previously uncharacterized 13 kD protein. Based on its structure and metal-binding properties, we hereby designate it FPM13 (*Francisella* Periplasmic Metalloprotein, 13 kDa). FPM13 is unique to the *Francisella* genus but highly conserved across species within the genus. In the complex, two donut-shaped nonameric rings stack together in a head-to-head fashion with D2 symmetry. Localization and biochemical assays confirmed that FPM13 is a soluble periplasmic protein capable of binding iron and copper. To gain insight into its biological function, we conducted extensive studies using *F. novicida* strains with clean in-frame deletions of the gene encoding FPM13. While its deletion does not impair bacterial growth in macrophages or mice, FPM13-deficient and epitope-tagged strains show increased copper sensitivity under iron-limited conditions, implicating a role in metal homeostasis and detoxification in *Francisella*.

## RESULTS

### Structural determination and identification of a novel *F. novicida* protein

The novel *F. novicida* protein was discovered in the process of characterizing a 150 kDa *Francisella* T6SS effector protein, PdpC. To determine the structure and function of PdpC, we expressed it with N-terminal FLAG and hexa-His epitope tags in *F. novicida*. A three-liter batch of bacteria expressing the epitope-tagged protein was pelleted by centrifugation, lysed by sonication, and the recombinant protein enriched by Nickel-agarose affinity chromatography followed by gel filtration on Sephacryl S200 (Fig. 1A). A Coomassie stained SDS-PAGE gel showed a dominant band corresponding to an ∼150 kDa protein, consistent with PdpC (Fig. 1B). However, a protein band at about 13 kDa which mostly eluted earlier on gel filtration (i.e., indicating a higher molecular weight complex) was also apparent on the Coomassie stained SDS-PAGE gel (Fig. 1B). Fractions from gel filtration with the most intense staining for the 150 kDa protein (Fig. 1B, Fractions 41 and 42) were analyzed by cryoEM (Suppl. Fig. S1A), with 2-dimensional class averages showing donut-shaped structures (Suppl. Fig. 1B). CryoEM analysis achieved 3.6 Å resolution (Suppl. Fig. S1C and Suppl. Fig. S2A), and the statistics are summarized in Table 1. The DeepTracer^11^ and cryoID program^12^ were subsequently run on the cryoEM density maps, using the *F. novicida* U112 genome as the search database to obtain a list of candidate proteins (Fig. 1C), leading to the identification of FPM13 (FTN_1118). Mass spectrometry analysis of the protein sample used for cryoEM analysis confirmed that PdpC is the dominant protein present and also revealed the presence of some other proteins (Fig. 1D), with FPM13 (FTN_1118) being the fifth most abundant protein in the sample used for cryoEM.

**Fig 1.**
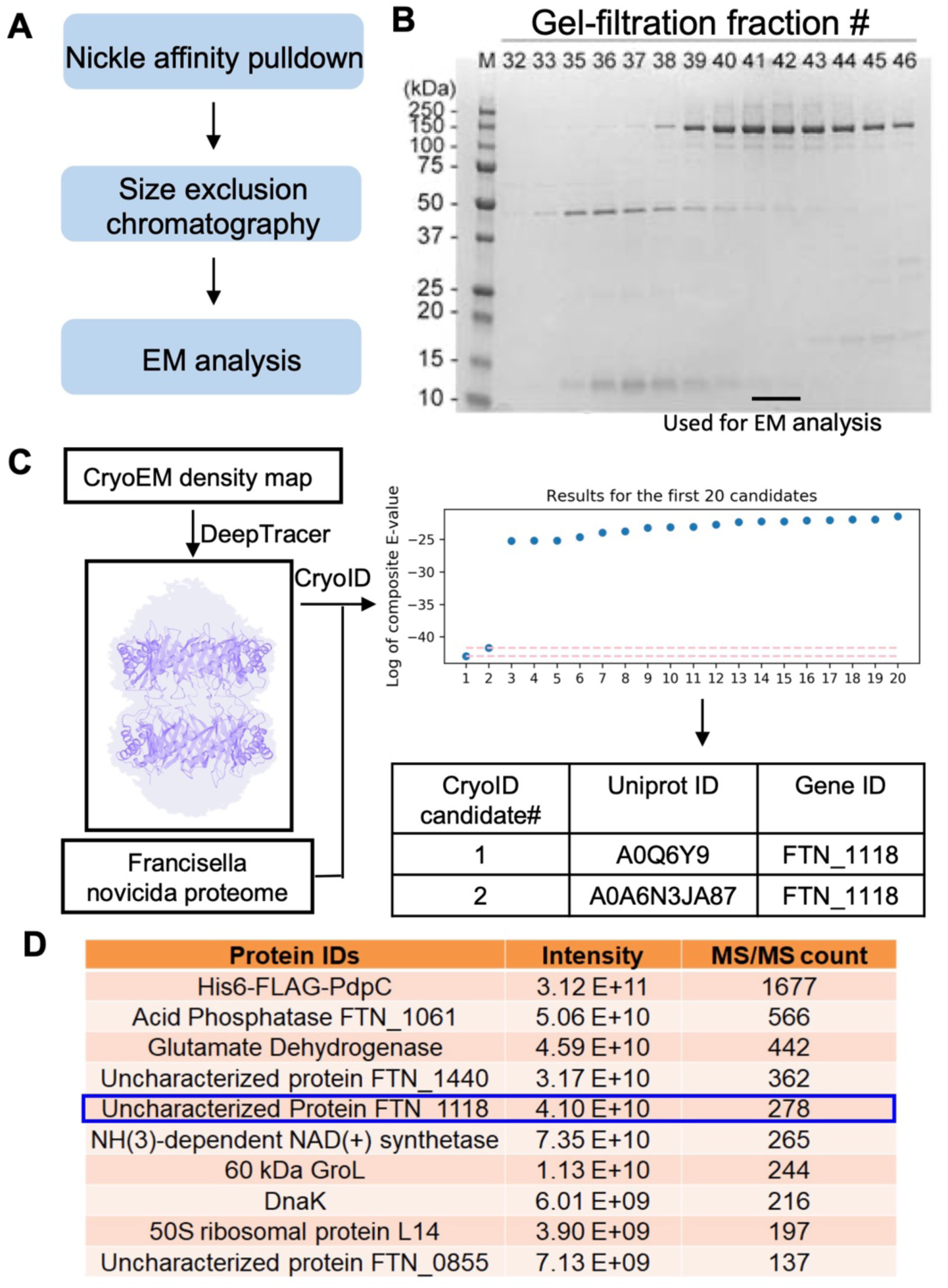
Discovery of FPM13 (FTN_1118) from *Francisella novicida.* **(A)** Workflow for protein sample preparation. **(B)** Coomassie blue stained SDS-PAGE gel for different fractions after gel-filtration. The fraction numbers are labeled. Fractions 41 and 42 are used for EM analysis. **(C)** Workflow for protein identification. The cryoEM map was used to generate a 3D model with predicted amino acid sequence traced by DeepTracer. The traced PDB model and the *Francisella novicida* proteome downloaded from UniProt are subjected to cryoID. The first ranking candidate is FTN_1118 (FPM13). De novo model building is performed in Coot. **(D)** The top10 proteins identified by mass spectrometry-based proteomics in the protein sample further verified the protein sample contains FTN_1118 (FPM13).

FPM13 is a protein composed of 111 amino acids. Amino acid residues 25 – 94 were resolved in the cryoEM density map (Fig. 2A), with the N- and C-termini (residues 1-24 and 95-111, respectively) being unstructured. The atomic model of the complex (Fig. 2B-D) shows a structure with two layers stacked head-to-head. Each layer has 9 repeating subunits of a monomer (13 kDa), consistent with the sequence of FPM13 and the ∼13 kDa band seen in the SDS-PAGE gels of the gel filtration samples (Fig. 1B). The entire structure has 18 repeats, corresponding to 234 kDa, explaining its high molecular elution profile on gel filtration. The complex measures ∼8 nm both in height and in outer diameter, with a 3.5 nm central channel (Fig. 2B), within which an uncharacterized density is observed (Suppl. Fig. S2B). It features a double-layered wall with an inner β-sheet core and an outer α-helical shell (Fig. 2C). Each monomer adopts a compact fold beginning with a β-strand, followed by an α-helix and two additional β strands (Fig. 3A-B). Comparison of the atomic model of the FPM13 monomer structure with structures in the Protein Data Bank did not yield any strong matches in a DALI^13^ search.

**Fig. 2:**
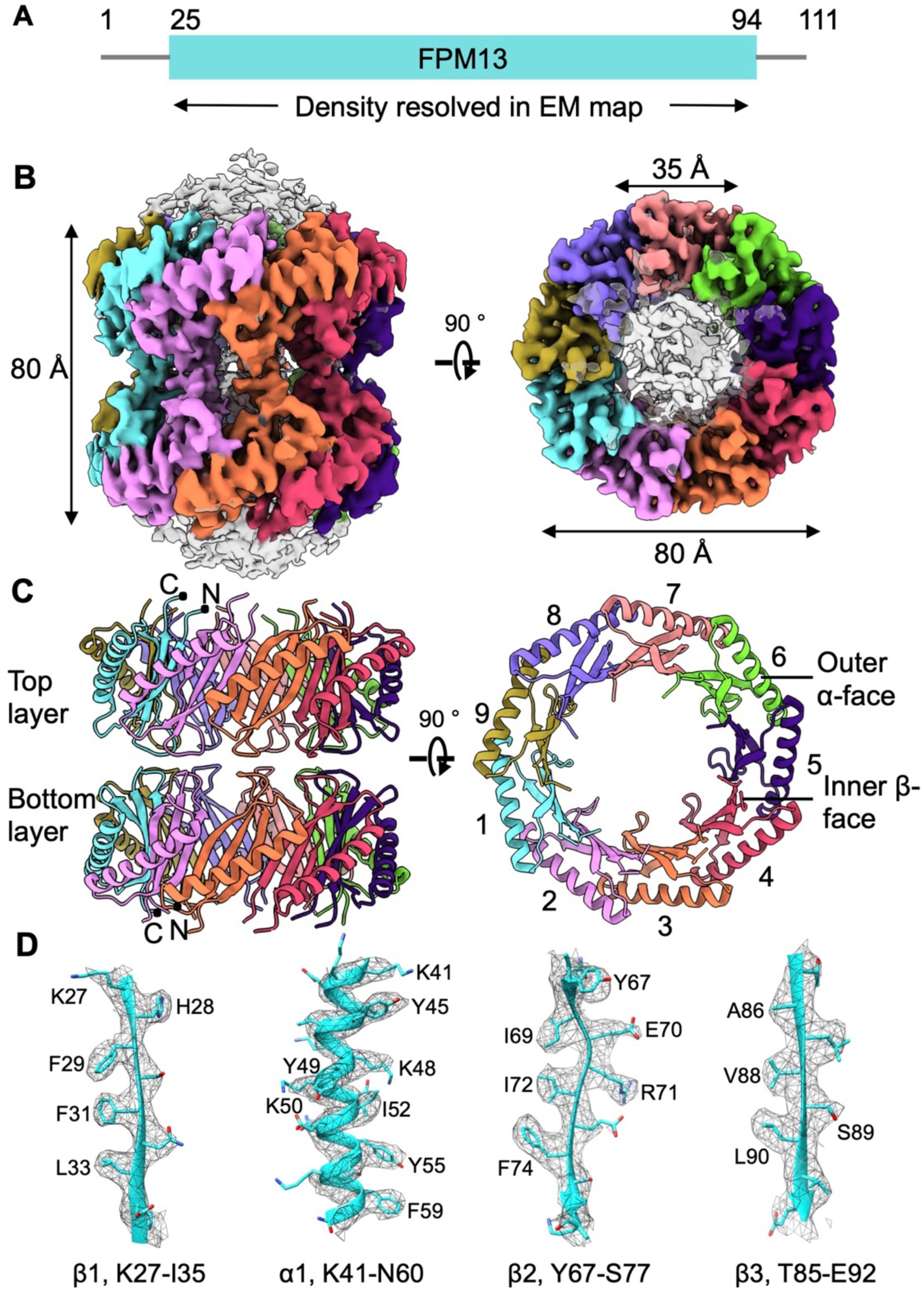
CryoEM structure of FPM13. **(A)** Diagram of *Francisella novicida* (Fn) protein FPM13. The full-length protein consists of 111 amino acids. S25-S94 are resolved in cryoEM map. **(B)** Two different views of the cryoEM density map of FPM13 octadecamer consisting of two nonamer layers stacked tail to tail. The protomers in each nonamer are represented in different colors. Two protomers stacked tail to tail within two layers are in the same color. **(C)** Two different views of FPM13 octadecamer structure. The structure is colored as in B. The protomers in one nonamer layer is labeled 1-9. **(D)** Representative density map of α helices and β strands.

**Fig 3.**
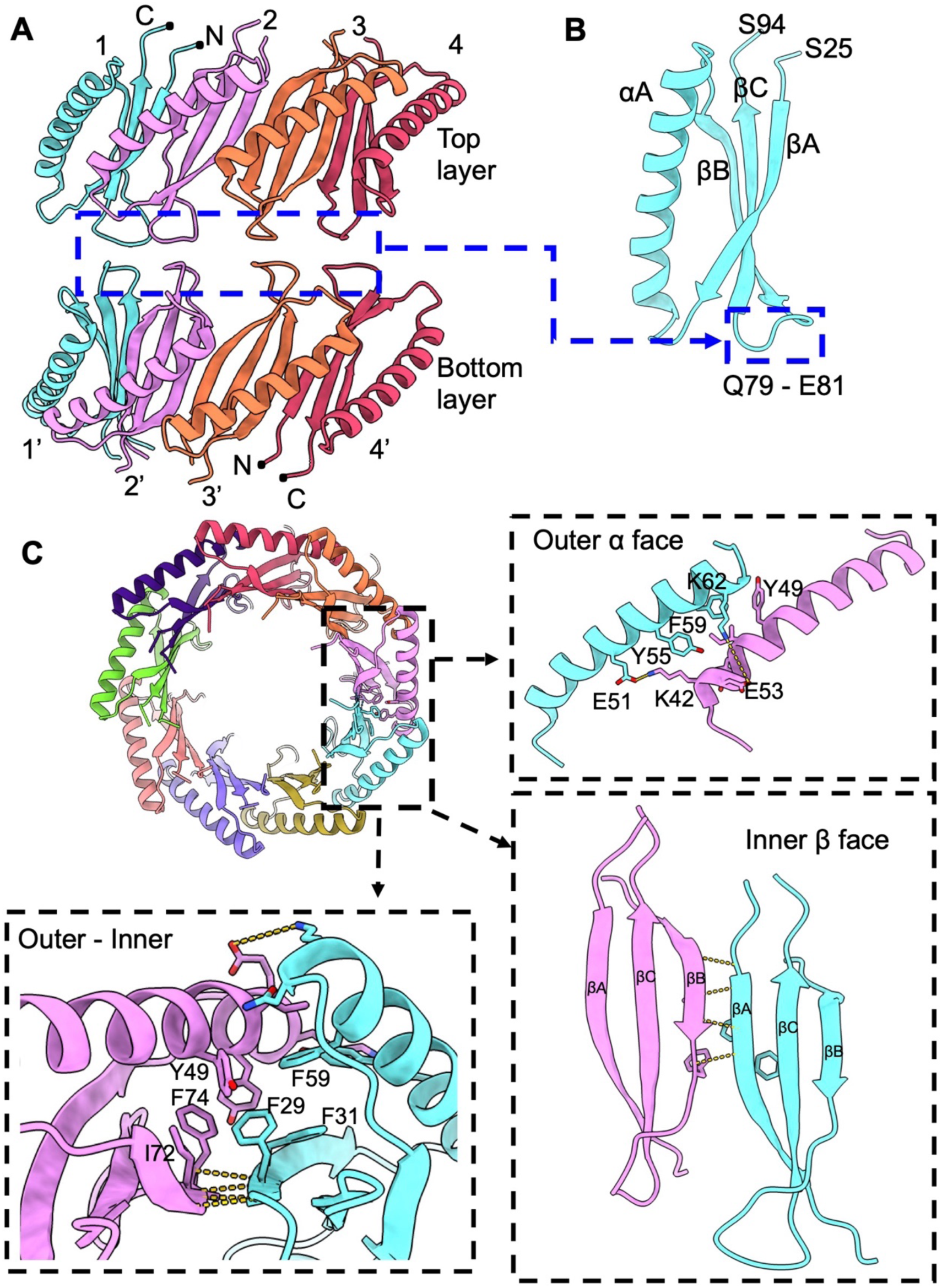
Interactions between FPM13 protomers. **(A)** Front view of the FPM13 assembly; the rear portion is omitted for clarity. **(B)** The loop region mediates the interactions between the top and bottom layers. **(C)** Illustration of the interactions between lateral subunits, with key residues shown in stick representation and labeled.

Interactions between FPM13 monomers in the assembly are shown in Fig. 3. The top and bottom layers of the 18-mer are stacked primarily through van der Waals interactions, mainly involving the loop regions spanning residues Q79 to E81 of the head-to-head protomers (Fig. 3A-3B). Lateral interactions between two subunits are mediated by a combination of hydrophobic (Y49, Y55 and F59) and electrostatic contacts (E51-K42, K62-E53) between the outer α helices (Fig. 3C). The inner β sheet is stabilized through backbone hydrogen bonding as well as hydrophobic interactions among side chains (Fig. 3C). The outer and inner faces are glued together by hydrophobic residues (Y49, F74, I72, F29, F31and F59). The interface between two adjacent subunits is substantial, with a buried surface area of 765 Å^2^, suggesting strong subunit association. We also confirmed the interaction of FPM13 with itself by bacterial 2-hybrid analysis (Suppl. Fig. S3).

### FPM13 is a soluble periplasmic protein

Bio-informatics analysis of the FPM13 sequence (PsortB^14^ and Uniprot^15^ annotation) indicated that the protein has a signal sequence and predicts that the protein is a periplasmic lipoprotein. To gain additional insights into the protein, we prepared *F. novicida* expressing FPM13 with a C-terminal ALFA epitope tag (FPM13-ALFA). We found that osmotic based preparations of *F. novicida* periplasmic proteins (e.g., obtained using the Tris-Sucrose-EDTA method) were contaminated with cytosolic proteins, such as GroEL and DnaK, thus complicating their interpretation. To determine whether FPM13 is a periplasmic protein, we utilized the technique of compartment-specific peroxidase-mediated biotinylation^16^ with APEX2, an engineered variant of soybean ascorbate peroxidase^17^. We prepared *F. novicida* that expresses FPM13-ALFA and the engineered ascorbate peroxidase either with (ssAPEX2) or without (APEX2) a signal sequence for secretion (Suppl. Fig. S4), thereby targeting the ascorbate peroxidase to either the periplasm or the cytoplasm, respectively. As shown in Fig. 4A, FPM13-ALFA was biotinylated and pulled down by streptavidin-agarose in the H_2_O_2_ treated *F. novicida* expressing ssAPEX2 targeted to the periplasm, but not in the strain expressing APEX2 lacking a signal sequence or in samples that were not treated with H_2_O_2_. As a positive control, the catalase-peroxidase enzyme, KatG (which has a signal sequence and is predicted to be periplasmic by PsortB) was also pulled down in the H_2_O_2_ treated ssAPEX condition (Fig. 4A). In contrast, the cytosolic protein GroEL was not pulled down in the ssAPEX conditions (with or without H_2_O_2_) but was pulled down in the *F. novicida* expressing cytosolically targeted APEX.

**Fig 4.**
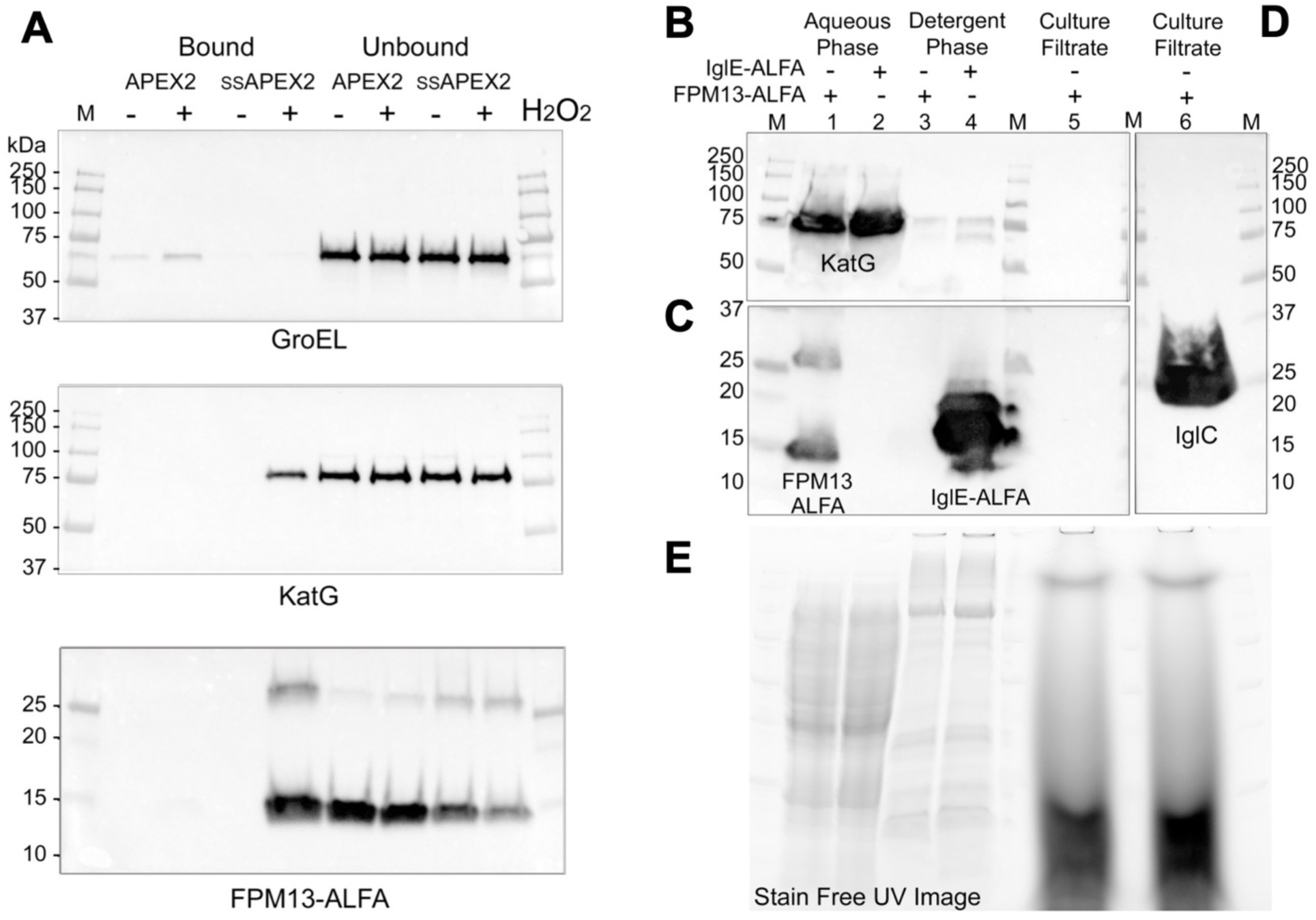
FPM13 is a soluble periplasmic protein. **(A)** FPM13 is biotinylated in the presence of H_2_O_2_ and pulled down by streptavidin-agarose APEX2 that is targeted to the periplasm (ssAPEX2), but not by cytosolic APEX2. GroEL and KatG serve as cytosolic and periplasmic control proteins, respectively. **(B - C)** TX-114 phase extraction shows that FPM13 and KatG are soluble proteins that partition into the aqueous phase, unlike the known lipoprotein, IglE (C, lane 4). FPM13-ALFA is not secreted into the culture filtrate (C, lane5), unlike positive control IglC (**D**, lane 6). Stain free UV-image shows equal loading of lanes being compared **(E).**

The ssAPEX2 biotinylation results indicated that FPM13 is located in the periplasm. To determine whether it is a lipoprotein, as indicated by PsortB and Uniprot annotation, we utilized the well-established method of TX-114 phase partitioning^18, 19^. As a control, we also prepared *F. novicida* expressing C-terminal ALFA-tagged IglE, a component of the T6SS apparatus that is well established as an outer membrane lipoprotein^19, 20^. We conducted TX-114 phase separation using the methods described by Brusca and Radolf^18^. While IglE-ALFA partitioned exclusively into the TX-114 phase, FPM13-ALFA partitioned exclusively into the soluble phase (Fig. 4C). As a control, the periplasmic protein KatG as expected partitioned into the aqueous phase (Fig. 4B). FPM13-ALFA was not detected in the culture filtrate (Fig. 4B), whereas the positive control, IglC, was detected strongly in the culture filtrate (Fig. 4D). Stain free UV image of the gel prior to transblotting confirmed comparable loading of the lanes being compared (Fig. 4E).

### FPM13 is not required for *F. novicida* growth in macrophages *in vitro* or in mice *in vivo*

FPM13 encodes a polypeptide of 111 amino acids in length and is of an unknown function. To determine whether FPM13 plays a role in *Francisella* growth in host cells, we generated a mutant strain of *F. novicida* U112 with deletion of the gene sequence corresponding to amino acid residues 1-92 (FnΔFPM13). We infected PMA-differentiated human THP-1 macrophages with the wildtype parent or FnΔFPM13 strain and harvested the bacteria from the monolayer at different time points after the infection (Fig. 5A). At 30 min post-infection, a similar number of wildtype and mutant strains were recovered from the monolayers indicating that there is no distinct difference in uptake of the two strains by host macrophages. After uptake, both the parent and mutant strains exhibited comparable growth kinetics with the bacterial number increasing by ∼1 log and 3 logs at 5 h and 22 h post-infection, respectively.

**Fig 5.**
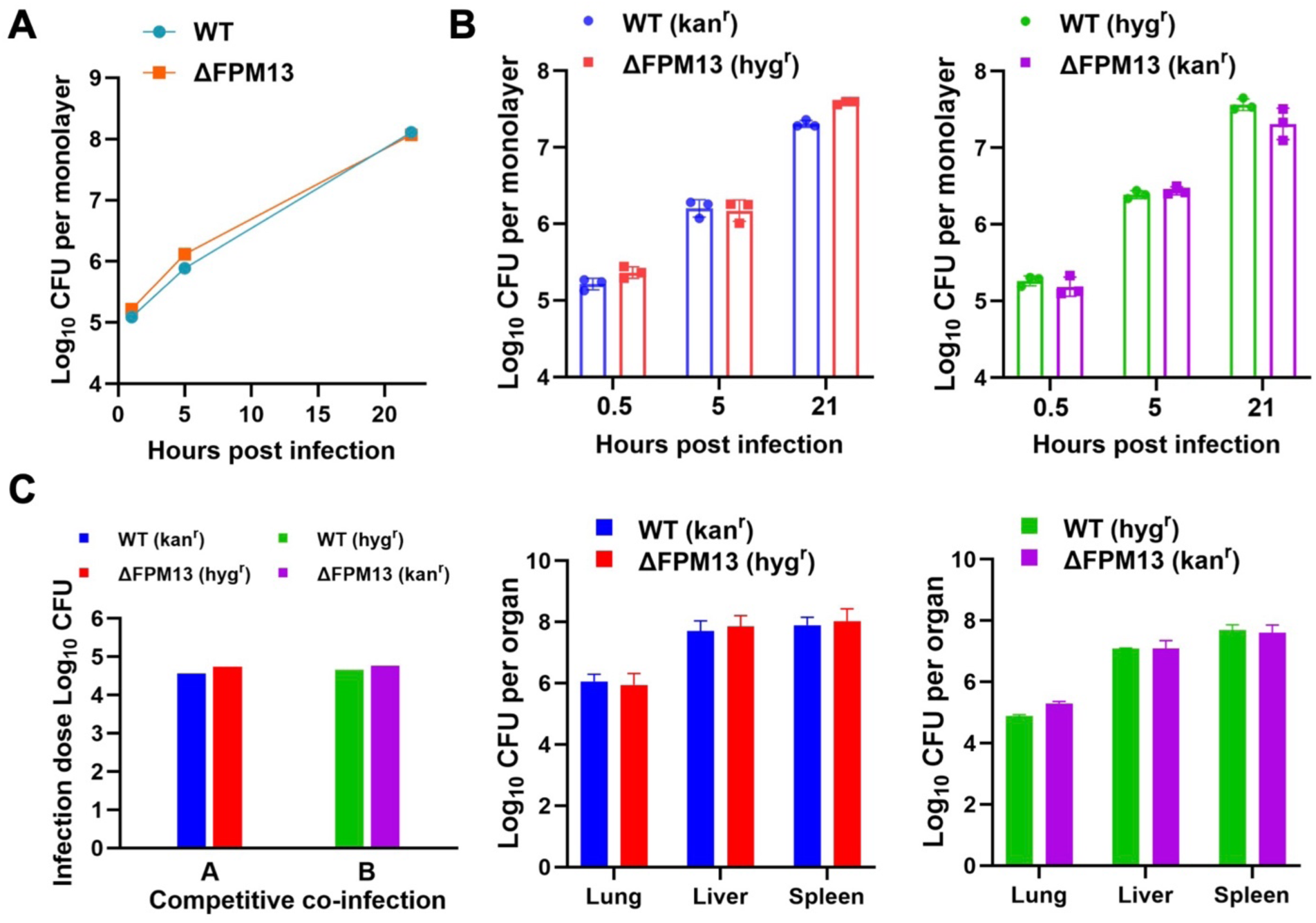
FPM13 is not required for growth in THP-1 macrophages in vitro or in mice in vivo. **(A)** PMA-differentiated THP-1 macrophages were infected with the wildtype parent (WT) or FnΔFPM13 strain of *F. novicida* and numbers of bacteria in macrophage monolayers determined by plating for CFU at the indicated time points. **(B)** Capacity of *F. novicida* WT and ΔFPM13 strains to compete for intracellular growth in THP-1 macrophages was examined in strains expressing kanamycin (kan) and hygromycin (hyg) resistance markers. Numbers of CFU of each strain in the monolayers was determined at the indicated times by plating monolayer lysates on agar containing kan or hyg. **(C)** Capacity of *F. novicida* WT and ΔFPM13 strains to spread to lung, liver and spleen after intraperitoneal infection was examined in a competition infection experiment using strains expressing hyg or kan resistance. Panel on the left shows the initial CFU dose of each strain used for infection. Middle and right panels show organ burdens of WT and ΔFPM13 strains with the indicated resistance markers.

Although FnΔFPM13 showed no obvious growth defect inside of THP-1 macrophages, it was conceivable that the deletion affects its fitness, a trait that often manifests under competitive growth of a mixed wildtype and mutant population in the same host environment. To examine this possibility, we introduced different antibiotic resistance markers (hygromycin and kanamycin) in the wildtype and mutant strains and assayed their ability to outgrow each other *in vitro* in THP-1 macrophages. To correct for any resistance marker associated bias, we performed the macrophage infection experiments with each of the resistance markers on either the wildtype or the deletion strain (Fig. 5B). We observed that the wildtype and mutant strains grow equally well in macrophages, with the strain bearing hygromycin resistance showing a slight competition advantage over the other strain bearing kanamycin resistance.

We next investigated the capability of the wildtype and FnΔFPM13 strains to establish an infection *in vivo* in mice. We pre-mixed the two strains at 1:1 ratio prior to infecting C57BL mice by the intraperitoneal route, and two days later we determined bacterial burdens in the lung, liver and spleen (Fig. 5C). The wildtype and Fn ΔFPM13 strains showed comparable growth in each of the organs suggesting that knocking out FPM13 has no impact on the fitness of the bacterium to spread and multiply in mice. In conclusion, our results showed that deletion of FPM13 has no discernible effect on the ability of *F. novicida* to replicate *in vitro* in macrophages or establish an infection *in vivo* in mice.

### FPM13 mutants exhibit an increased sensitivity to CuSO_4_

We observed no difference between FPM13 mutants and wild-type *F. novicida* in sensitivity to H_2_O_2_, erythromycin, ethanol, polymyxin B, or osmotic shock. However, we found that in strains of *F. novicida* that were depleted of iron and grown with high aeration (shaking at 200 rpm), bacteria that expressed C-terminally tagged FPM13 and those in which FPM13 was deleted are much more sensitive to CuSO_4_ (Fig. 6). We did not observe this phenotype in bacteria that were not iron depleted or in bacteria that were grown without shaking (i.e., in stationary 96-well plates). We hypothesize that FPM13 may be important in metal ion transport or detoxification. Copper exerts its toxicity by mis-metalation of proteins and by increased formation of reactive oxygen species. Both mis-metalation and formation of reactive oxygen species would be expected to be more pronounced in bacteria that have been depleted of iron and grown with high aeration, thereby favoring observation of the FPM13 copper-related phenotype.

**Fig 6.**
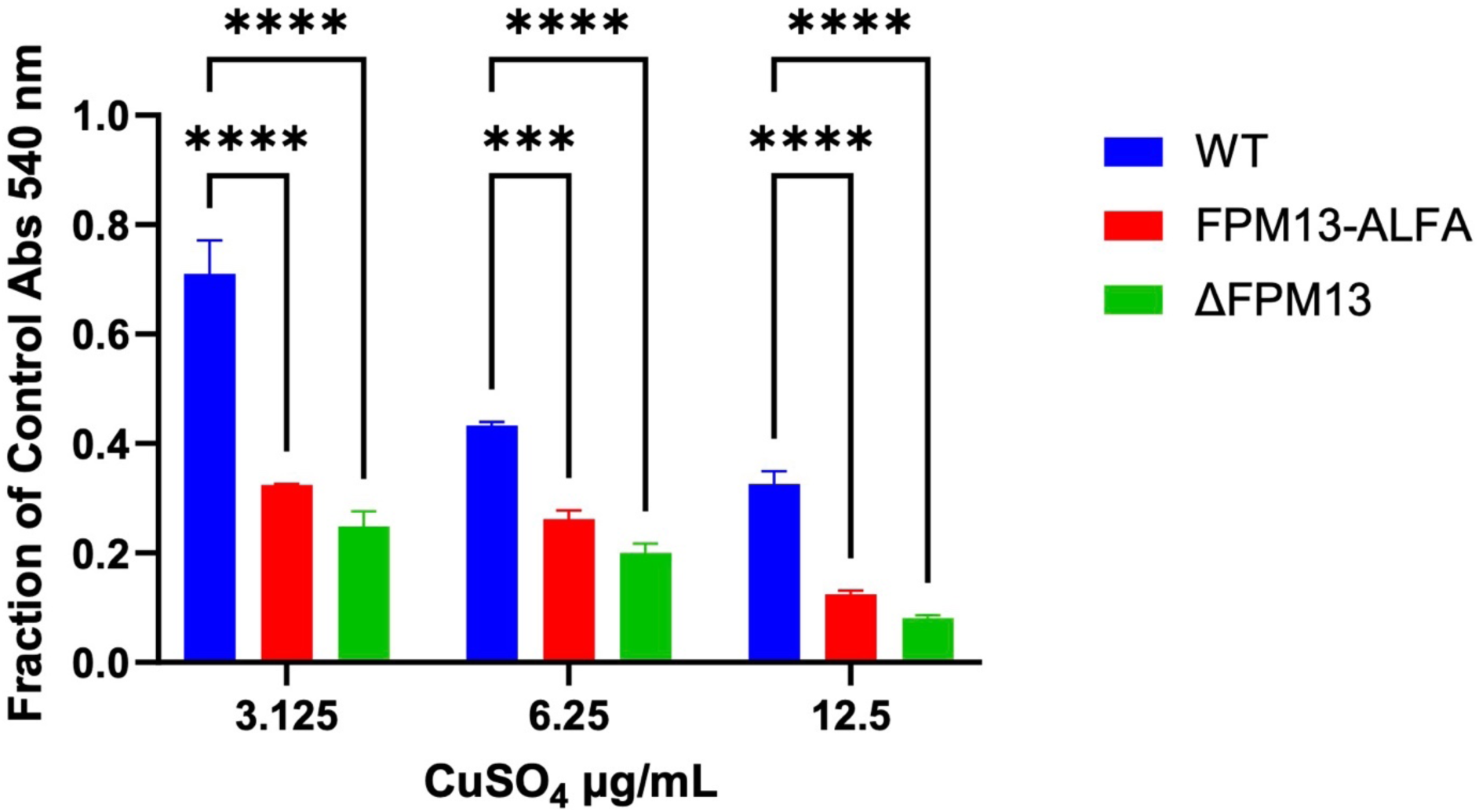
*F. novicida* FPM13 mutants that have been depleted of iron show increased sensitivity to CuSO_4_ compared with the wild-type strain when grown with high aeration. Data indicate mean ± standard error of independent triplicate determinations. Statistical analysis by two-way Anova with Dunnett’s correction for multiple comparisons. *** *p* < 0.001; **** *p* < 0.0001.

### FPM13 homologs are restricted to *Francisella* Clade A

Based on a phylogenetic analysis of 425 orthologous single-copy genes, Duron et al.^21^ divided *Francisella* into three clades: one corresponding to mammalian pathogens and terrestrial environments (including *F. tularensis* and *F. novicida* species); one corresponding to fish pathogens (including *F. philomiragia* and *F. noatunensis*); and a separate distinct clade of tick endosymbionts (including *F. persica*). Kuman et al.^22^ performed a pan-genome analysis of 63 *Francisella* genomes and achieved a similar categorization, identifying two distinct clades: Clade A corresponding to *F. tularensis* and *F. novicida* species (species mainly associated with terrestrial animals) and Clade B (*F. philomiragia* and *F. noatunensis*, species associated with aquatic animals). Their analysis also identified a third more diverse cluster, designated Clade C, found mainly in marine environments and comprising *Allofrancisella guangzhouensis* 08HL01032T, *F. frigiditurris* sp. Nov. CA971460, *F. endociliophora* FSC1006, *F. uliginis* sp. nov. TX077310, and *F. halioticida* DSM23729. An additional four outlier species (*F. hispaniensis* FSC454, *F. cf. novicida* 3523, *F. opportunistic asp*. Nov. MA067296, and *F. persica* ATCC VR331) reside between Clades A and B, but closer to Clade A^22^.

The FPM13 sequence is unique to *Francisella,* as BLAST^23^ homology searches do not identify any homologs outside of *Francisella*. We have found that the FPM13 protein sequence is highly conserved among *Francisellaceae* and homologs are present in all Clade A species of *Francisella* (including *F. holarctica, F. mediasiatica, F. tularensis,* and *F. novicida*) as indicated in Suppl. Table S2. However, it is absent from the tick endosymbionts, including *F. persica*, and it is absent from all Clade B species (fish pathogens and marine isolates *F. philomiragia*, *F. saliminara*, and *F. noatuensis*). FPM13 homologs are also absent from almost all Clade C species, including the fish pathogen *F. halioticida*, the ciliate pathogen *F. endociliophora,* and species isolated from water samples such as *A. guangzhouensis* and *F. uliginis*. Intriguingly, an FPM13 homolog is present in the Clade C member *F. frigiditurris*, which was isolated from the water of an air conditioning system. Further comparisons of *Francisella* species that do and do not possess FPM13 may help to reveal its biological function.

### FPM13 is a metalloprotein

We noticed that FPM13-ALFA purified by affinity chromatography with anti-ALFA resin was colored, with an absorbance maximum of 410 nm (Fig. 7A-C), suggesting that it is a metalloprotein. ICP-MS results confirmed that the protein contains both Fe and Cu in metal:octadecamer molar ratios of 4.2 and 3.3, respectively (Fig. 7E). As the protein was purified in the presence of 1 mM EDTA, it is possible that higher metal:protein ratios are possible. We initially discovered FPM13 by virtue of its binding to Ni-agarose and elution in high molecular weight fractions on gel filtration (Fig. 1). The binding to Ni-agarose, and potentially other metals, is explained by the presence of cysteine and histidine residues in positions well suited for co-ordination of metal ions (Fig. 7D and Suppl. Fig. S5).

**Fig 7.**
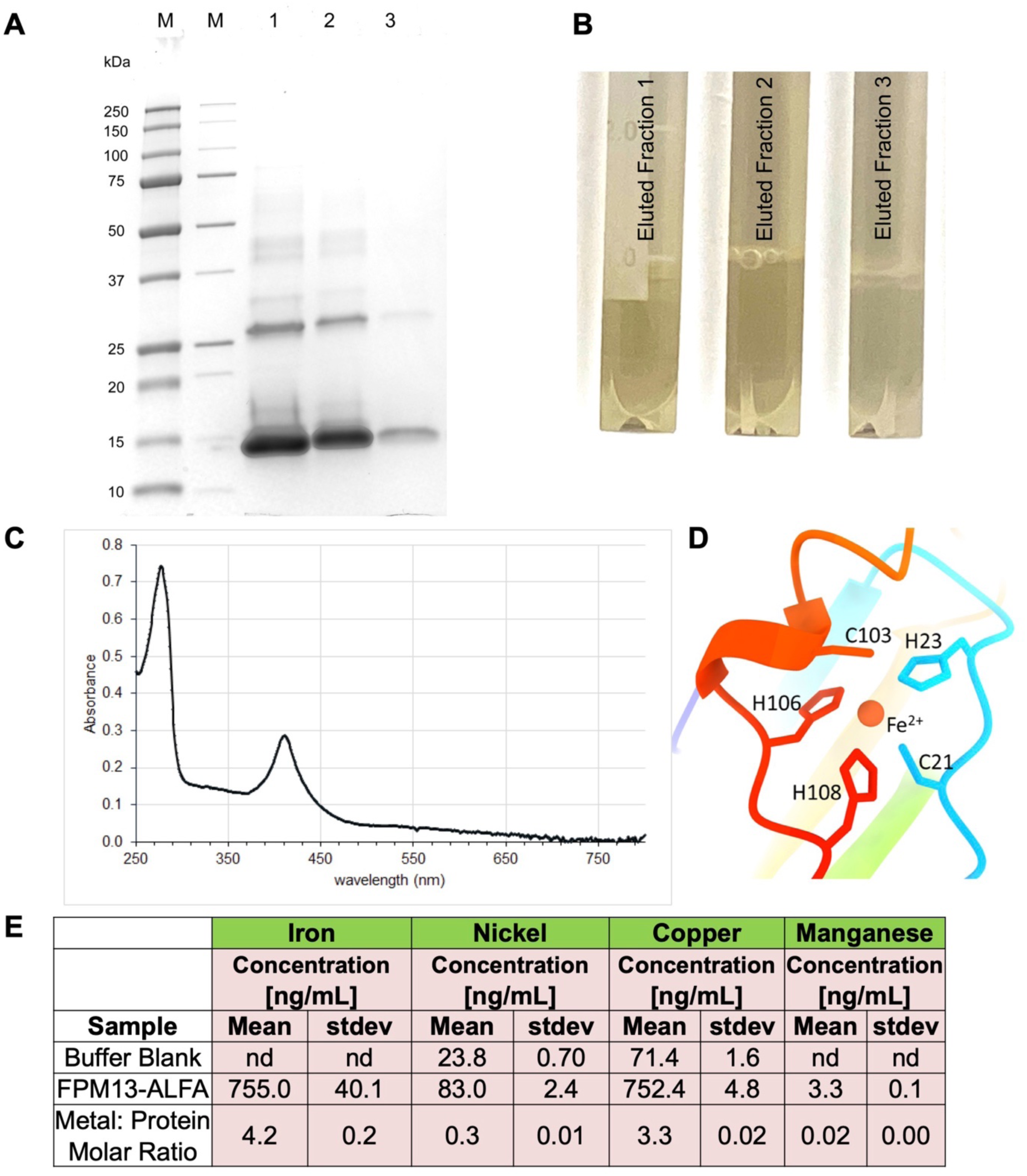
Identification of FPM13 as a metalloprotein. **(A)** Coomassie stained gel showing purification of FPM13-ALFA on ALFA-select resin. Lanes 1 - 3 correspond to consecutive elutions of resin with 0.1 M glycine HCl, pH 2.2. **(B)** Appearance of eluates 1 – 3 demonstrating brown color. **(C)** UV-vis spectrum of fraction #1, showing absorbance peaks at 280 nm and 410 nm. **(D)** Ribbon diagram illustrates possible mechanism of metal binding predicted by Chai-1. **(E)** ICP-MS showing presence of Cu and Fe in the sample eluted from the resin. Data from a buffer blank is also shown. Data represent means of triplicate determinations. “nd” indicates below the detection threshold. Metal to FPM13 octadecamer molar ratios are indicated based on a measurement of 0.87 mg/mL protein in the sample used for ICP-MS.

## DISCUSSION

We have identified FPM13, a novel 18-mer periplasmic protein from *F. novicida*. Deleting the protein or adding a C-terminal epitope tag to the protein leads to increased sensitivity to CuSO_4_ in bacteria that have been depleted of iron. We hypothesize that FPM13 and its homologs in other terrestrial *Francisella* is involved in transport or detoxification of metals. It is likely that addition of the C-terminal epitope tag disrupts the interaction of FPM13 with proteins required for metal transport or detoxification such that – like the FPM13 deletion strain – it shows increased sensitivity to CuSO_4_. The protein is highly conserved in *Francisella* species that are found in terrestrial environments (*F. tularensis* and *F. novicida* species), but it is absent from *Francisella* that are fish pathogens (*F. philomiragia*, *F. noatunensis*, *F. halioticida*). It has been proposed that macrophages^24^ and possibly amoebae^25^ use elevated phagosomal Cu and/or Zn intoxication as part of an innate immune response (“brass dagger”) to combat intracellular bacterial pathogens^24^. The *Francisella* tick endosymbionts do not face such innate immune responses in their hosts, and thus may not require the metal detoxification and transport machinery required by pathogenic bacteria, explaining why they lack FPM13 homologs. *Francisella* of Clades B and C may have evolved different methods for detoxifying and transporting metals in their quite different aquatic and marine environments. It is likely that *Francisella* have redundant pathways for transporting and detoxification of metals, accounting for why we did not observe a phenotype in our macrophage or mouse infection studies. Additional research is needed to identify the protein machinery that interacts with FPM13 for metal transport or detoxification.

This study underscores the transformative potential of the cryoID approach in structural biology, offering significant advantages over traditional methods reliant on protein expression with affinity tags. Unlike conventional approaches, which require the target protein to be tagged, overexpressed, and purified prior to structural analysis, cryoID enables de novo identification of unknown proteins directly from cryoEM density maps. This capability facilitated the discovery of FPM13, a novel *Francisella* periplasmic metalloprotein, without prior knowledge of its sequence or function. By bypassing the need for recombinant expression, cryoID preserves native protein conformations, capturing less populous, transiently stable states, as well as interactions with cofactors, metal ions, and binding partners that might be disrupted during purification. Furthermore, cryoID streamlines structural determination by reducing the reliance on labor-intensive cloning and purification steps, accelerating the study of complex macromolecular assemblies. These advantages position cryoID as a powerful tool for uncovering novel proteins and elucidating their biological roles in their native cellular contexts, with broad implications for advancing structural biology and proteomics research.

## METHODS

### Generation of *F. novicida* strains

Allelic exchange cassettes for in-frame deletion of FPM13 (ΔFPM) and replacement with a C-terminally ALFA-tagged FPM13 (FPM13) including its 1.2 kb genomic upstream and downstream sequences were cloned into pMP590 using standard restriction cloning procedures^26^. The plasmid constructs were confirmed with nucleotide sequencing (Primordium) prior to introducing into *F. novicida* U112. Transformants grown on chocolate GCII agar containing 15 μg/ml kanamycin were spotted on chocolate GCII agar without antibiotics and the presence of the allelic exchange cassette was confirmed using colony PCR. Counterselection using the sacB marker on pMP590 was performed by streaking transformants on chocolate GCII agar containing 7% (w/v) sucrose. Strains that grew on CA-sucrose and were sensitive to kanamycin were screened using colony PCR to confirm gene deletion or replacement. Chromosomal deletion was validated by PCR amplification of the targeted genomic region and sequencing of the PCR amplicon. Replacement with epitope tagged FPM13 was validated by Western blotting and detecting with ALFA-HRP (Synaptic Systems, 1:4000) antibodies.

APEX2 sequence was amplified by PCR from Vimentin - APEX2 in pECFP, a gift from Alice Ting^27^ (Addgene plasmid #66170). The expression cassette of 3xFLAG-tagged APEX2 with or without the signal peptide sequence from *F. novicida* beta-lactamase (BlaB) driven by the promoter of bacterioferritin from *F. tularensis* Live Vaccine Strain was inserted into the *blaB* locus in the *F. novicida* strain expressing FPM13-ALFA to replace a genomic region from 30 nucleotides upstream to the first 801 nucleotides of the *blaB* gene.

Plasmids pMP607 and pMP633 bearing a kanamycin or hygromycin resistance marker, respectively,^26^ were transformed into *F. novicida* U112 with and without FPM13 deletion for assaying competitive growth *in vitro* in macrophages and *in vivo* in mice.

### *F. novicida* intracellular growth assay in THP-1 cells

Human monocytic THP-1 cells were maintained in advance RPMI medium supplemented with 2% heat-inactivated fetal bovine serum (HI-FBS), glutamax, and penicillin/streptomycin. The cells were differentiated with phorbol 12-myristate 13-acetate in antibiotic-free medium with 10% HI-FBS for 3 days and infected with overnight cultures of *F. novicida* strains at a ratio of 1:1 (bacterium:cell) at 37°C for 90 min. The infection medium was replaced with medium containing 10 μg/ml gentamicin and incubated at 37°C for 30 min after which the monolayer was washed twice and supplied with fresh culture medium containing 0.2 μg/ml gentamicin. At various time points after infection, monolayers were lysed with 1% saponin in phosphate buffered saline (PBS) at room temperature for 5 min, and the lysates were serial diluted in PBS and spotted on plates for colony-forming units (CFU). For competitive growth assay, equal numbers of *F. novicida* strains bearing different antibiotic resistance markers were premixed prior to infecting PMA-differentiated THP-1 cells. At indicated times, bacteria were harvested from infected monolayers and CFU enumerated after spotting on both agar plates containing kanamycin and agar plates containing hygromycin.

### Competitive growth of *F. novicida* strains in mice

Female C57BL6 mice aged of 7-10 weeks (Jackson Laboratory) were injected intraperitoneally with 0.1 ml normal saline (Cytiva HyClone) containing a mixture of 1×10^5^ CFU each of the *F. novicida* U112 parental and ΔFPM13 strains, carrying kanamycin or hygromycin resistance markers, respectively, or vice versa. Two days later, their lungs, liver, and spleen were harvested and homogenized in PBS. Serial diluted homogenates were plated on GCII chocolate agar containing 10% IsoVitalex and 15 μg/ml of kanamycin or 200 μg/ml hygromycin for enumeration of CFU.

### Bacterial 2-Hybrid protein interaction assay

*F. novicida* gene encoding FPM13 was cloned into each of the four expression vectors (pUT18, pUT18C, pKNT25, and pKT25) of a bacterial adenylate cyclase-based two-hybrid (BACTH) system (Euromedex) in-frame with the coding sequence of two complementary T18 or T25 fragments of *Bordetella pertussis* adenylate cyclase (CyaA). After confirmation by nucleotide sequencing, a pair of compatible plasmids for expression of FPM13 with N- or C-terminal T18 and T25 fusions were co-transformed into the *cya*A negative *E. coli* strain BTH101 by electroporation. Transformants were streaked on Lurie agar containing kanamycin (50 μg/ml), carbenicillin (100 μg/ml), IPTG (1 mM), and X-gal (40 μg/ml), and the plates were incubated at 30°C for two days.

### Protein purification

PdpC: *Francisella novicida* expressing FLAG- and His_6_-epitope tagged PdpC was grown in trypticase soy broth with 0.2% cysteine (TSBC), 5% KCl, 0.1 mg/mL FeSO_4_, and 5 mM betaine to an optical density (540 nm) of 2.0. Bacteria were pelleted by centrifugation (4000 g for 90 min at 4°C), resuspended in lysis buffer (50 mM sodium phosphate, pH 7.4, 1 mM EDTA, 1 mM PMSF, 1 mM NEM, and 1:100 protease inhibitor cocktail (HY-K0010, MedChemExpress), and lysozyme (1 mg/mL)) and disrupted by sonication on ice with a probe tip sonicator. The sonicate was clarified by ultracentrifugation (44,400 g for 90 min at 4°C) and imidazole, NaCl, and Mega-9 detergent added to achieve concentrations of 10 mM, 0.15 M and 0.2%, respectively. The supernate was rotated overnight with 0.5 mL of high capacity, EDTA compatible, nickel-agarose (Pierce), washed extensively with 50 mM sodium phosphate with 0.15 M NaCl, 1 mM EDTA, 10 mM imidazole, and eluted with an imidazole step gradient (20 mM, 50 mM, 100 mM, 250 mM). Eluted fractions were analyzed by SDS-PAGE and those containing a 150 kDa band corresponding to PdpC were concentrated with a 100 kDa MWCO Centrifugal filter (Amicon Ultra 15), applied to a Sephacryl S200 gel filtration column and eluted in 50 mM sodium phosphate pH 7.4, 0.15 M NaCl, 1 mM EDTA, 0.2% Mega-9. Fractions were analyzed by SDS-PAGE and the fractions corresponding to the peak of the 150 kDa band were concentrated with a 100 kDa MWCO centrifugal filter and used for cryoEM.

FPM13: *F. novicida* expressing FPM13-ALFA was grown in in 3 liters of TSBC to an optical density (540 nm) of 2.0. Bacteria were pelleted, resuspended in lysis buffer, sonicated, and clarified by ultracentrifugation as described above. The clarified supernate was added to 0.5 mL of ALFA-select resin CE (Synaptic Systems) and rotated overnight at 4°C. The resin was washed with 20 mM sodium phosphate, pH 8.0, 150 mM NaCl, 0.5% TX-100, followed by washing in 20 mM sodium phosphate, pH 8.0, 150 mM NaCl. The washed resin was transferred to a column and bound protein was eluted with consecutive applications of 0.8 mL of 0.1 M glycine HCl, pH 2.2, 0.8% CHAPS with immediate neutralization by collection of the eluate in tubes containing 0.2 mL of 1 M Tris HCl, pH 8.5. The sample eluted from the column was analyzed by SDS-PAGE, UV-Vis spectroscopy, and inductively Coupled Plasma-Mass Spectrometry (ICP-MS).

### CryoEM sample preparation and image collection

The graphene grids were prepared similarly to as described in the previous work^28^ and were used on the same day. An aliquot of 3 μL of purified protein sample was applied to prepared graphene grids. After incubation for 30 s, the grid was blotted for 8 s with 8 blotting force at 4°C and 100% humidity in an FEI Vitrobot Mark IV (Thermo Fisher Scientific). The grid was then flash-frozen in liquid ethane and stored in liquid nitrogen for data collection.

Images were collected using a Titan Krios electron microscope (Thermo Fisher Scientific) at 300 kV. The microscope was operated with the GIF energy-filtering slit width setting to 20 eV in super-resolution mode. Movies were acquired with SerialEM^29^ at a magnification of ×81,000, corresponding to a pixel size of 1.1 Å on the sample level, with exposure time of 2 s and total dosage of ∼50 electrons/Å2, dose-fractionated into 40 frames. 19,280 movies were collected.

### Single-particle cryoEM reconstruction

The workflow for cryoEM reconstruction is summarized in Supplementary Fig. S2. Frames of each movie were aligned for correction of beam-induced drift with MotionCor2^30^, generating two averaged images, one with dose weighting (used for particle extraction and further reconstructions) and the other without (for contrast transfer function determination, particles picking and defocus determination). The pixel size of averaged images is 1.1 Å on the specimen scale. The defocus values of images were determined by CTFFIND4^31^. The following data processing was done with RELION 3.1^32, 33^. 22,936,178 particles were automatically picked, extracted in dimensions of 2×-binned to 90 × 90 pixels (2.2 Å per pixel) to speed up the data processing procedure. 9,453,347 particles were selected from 2D classification. 3D classification was followed, and 3,040,320 particles were selected. Then the selected particles were re-extracted with a box size dimension of 180 × 180 pixels (1.1 Å per pixel) and subjected to local-Refinement. The particles after refinement were further re-extracted with a box size dimension of 300 × 300 pixels (1.1 Å per pixel), followed by local-Refinement. Skip align class3D was applied and two good classes were selected and combined. Another 3D refinement was applied, generating a final 3.6 Å-resolution reconstruction. The resolution of the map was estimated from the gold-standard Fourier shell correlation criterion, FSC = 0.143. Data collection and processing statistics are summarized in Table S1.

### Protein identification and model building

The reconstructed cryoEM map was used to generate a 3D model trace using DeepTracer^11^. The sequence model provided by DeepTracer was then analyzed with cryoID^12^ against the *Francisella novicida* protein database in Uniprot. The top-scoring candidate was identified as FTN_1118 (FPM13). The atomic model of FPM13 was built and manually adjusted in COOT^34^. The model was then refined using Phenix^35^ in real space with secondary structure, Ramachandran and rotamer restraints. Refinement statistics of the model were given in Table S1. Figures and movies were generated with UCSF Chimera^36^ and Chimera X^37^.

### ICP-MS

ICP-MS analysis was conducted to identify and quantitate metal ions in the sample eluted from the ALFA-resin select CE and a buffer control. The samples were transferred to clean Teflon vessels and digestion was carried out with concentrated HNO_3_ (65-70%, Trace Metal Grade, Fisher Scientific) with a supplement of H_2_O_2_ (30%, Certified ACS, Fisher Scientific) at room temperature for 2 hours. After complete digestion of the sample, it was diluted to a final volume of 5 mL by adding filtered deionized water for analysis. The calibration curve was established using a standard solution while the dwell time was 50 ms with thirty sweeps and three replicates with background correction.

### Mass spectrometry

Protein in the sample used for CryoEM analysis was acetone precipitated by adding 4 volumes of acetone, storing the samples overnight at -20°C, and centrifuging at 10,000 g for 10 min at 4°C. The pellet was washed twice with 4:1 acetone:water at 4°C, air dried for 30 min at room temperature, and stored at -20 °C. Further processing was conducted by the UCLA Proteome Research Center. The pellet was resuspended in 8 M urea, 100 mM Tris-HCl, pH 8.5; reduced with 5 mM tris(2-carboxyethyl)phosphine (TCEP); alkylated with 10 mM iodoacetamide; and digested with sequencing-grade trypsin. The peptide mixture was desalted, fractionated on-line using C18 reversed phase chromatography, and analyzed using tandem mass spectrometry on a Q-Exactive mass spectrometer (Thermo Fisher Scientific). Data analysis was performed using IP2 (Integrated Proteomics Applications) against a Fn U112 database (taxid: 401614) and filtered using a decoy-database estimated false discovery rate of less than 0.01.

### Western blotting

Proteins were separated using Any kD Mini-Protean TGX Stain-free gels (Bio-Rad) and visualized using a ChemiDoc Imaging System (Bio-Rad) prior to transblotting onto a 0.2 μm nitrocellulose membrane. The membrane was blocked with EveryBlock blocking buffer and probed with HRP-conjugated camelid single domain antibody to ALFA epitope tag (Synaptic Systems) or rabbit antibody to KatG or GroEL^38^ at a dilution of 1:4000 (anti-ALFA) or 1:1000 (rabbit primary antibodies), respectively. HRP-conjugated goat anti-rabbit antibody (Bio-Rad) was used as a secondary antibody and chemiluminescent signals were developed by incubating with Clarity Western ECL substrates (Bio-Rad) and detected using the ChemiDoc Imaging System.

### APEX2 labeling

The bacteria were grown in trypticase soy broth to an optical density of 1.8 and incubated with 1 mM biotin-phenol (Cayman Chemical) for one hour prior to addition of 1 mM H_2_O_2_ (Sigma Chemical Company). Cultures processed identically but without addition of H_2_O_2_ were prepared as controls. After 3 minutes, the *in vivo* biotinylation reaction was terminated by addition of sodium azide, sodium ascorbate, and Trolox to achieve final concentrations of 10 mM, 10 mM, and 5 mM, respectively. The bacteria were pelleted by centrifugation (10,000 g for 10 min) and washed twice with TBS containing 10 mM sodium azide, 5 mM Trolox and 10 mM sodium ascorbate. The washed bacterial pellets were resuspended in RIPA buffer (50 mM Tris HCl, pH 8, 0.15 M NaCl, 1% Nonidet P-40, 0.1% SDS, 0.05% sodium deoxycholate, 1.5 mL for each 7.5 mL of original culture) and sonicated on ice with a probe tip sonicator. Insoluble material was pelleted by centrifugation at 10,000 *g* for 60 min at 4°C and the soluble supernate was incubated overnight with streptavidin-agarose (Novagen, 50 uL of washed resin per 1.5 mL clarified supernate). The resin was washed with RIPA buffer and bound biotinylated proteins were eluted by heating to 90°C for ten minutes in SDS-PAGE sample buffer.

### TX-114 phase extraction

*F. novicida* expressing FPM13-ALFA or IglE-ALFA were grown to an optical density of 2.0 in TSBC. PEG-10,000 (10%) was added to the IglE-ALFA culture to induce T6SS expression. Bacteria were pelleted by centrifugation at 10,000 g for 10 min, resuspended in TBS containing 1 mM EDTA, 1 mM PMSF, 1 mM NEM, and 2% TX-114 (2 mg bacterial protein/mL). The culture supernate was saved for analysis of secreted proteins. Suspensions were sonicated on ice with a probe tip sonicator, stirred for one hour in an ice bath, and insoluble material removed by centrifugation at 10,000 g for 1 hour at 4°C. The samples were incubated at 37°C for 10 min to induce phase separation and phases were separated by centrifugation at 20,000 g for 15 min at 23°C. The aqueous (upper) phase was removed and 10% TX-114 added to achieve 2% concentration. Four volumes of TBS containing 0.04% TX-114 was added to the lower detergent phase. The samples were mixed at 4°C for one hour and phase partitioning repeated as before. The aqueous wash and detergent wash of the detergent and aqueous phases, respectively, were discarded and the washing process was repeated twice. Protein of the final washed aqueous and detergent phases and of the culture supernate was precipitated by addition of ten volumes of acetone (-30°C). After 18 hours at -30°C, the precipitated protein was pelleted by centrifugation at 20,000 g for 10 minutes, dried briefly, and resuspended in SDS-PAGE sample buffer for analysis by Western Immunoblotting.

### Copper sensitivity

*F. novicida* strains (WT, FPM13-ALFA, and Λ1FPM13) were inoculated into Trypticase Soy Broth with Cysteine (TSBC) from frozen stocks and grown overnight to an optical density of 1.8 – 2.2. The bacteria were pelleted by centrifugation at 22°C, washed three times with PBS, and inoculated into Chamberlain’s Chemically Defined Medium^39^ (CDM) without any added iron at an optical density (540 nm) of 0.05. The bacteria were depleted of iron by growth in this iron deficient medium at 37°C for 4 - 6 hours until they reached an optical density of 0.5 (we did not observe any difference in rate of growth). The iron depleted bacteria were inoculated into CDM with 0.67 µg/mL FeSO_4_ with 0 – 25 µg/mL CuSO_4_. Cultures of 0.5 mL in 5 mL Falcon 12 x 75 mm tubes (Corning) were incubated at 37°C, shaking at 200 rpm for 18 hours and optical densities were measured by spectrophotometer at 540 nm.

## Acknowledgements

This project was supported by National Institutes of Health grant AI151055 (MAH and ZHZ). The authors thank Chong Hyun Chang for assistance in performing the ICP-MS, which was conducted in the ICP-MS facility within the Nano and Pico Characterization Laboratory in CNSI at UCLA. We acknowledge the use of resources at the Electron Imaging Center for NanoSystems supported by US NIH (1S10OD018111) and the US National Science Foundation (DBI-1338135 and DMR-1548924).

## Author contributions

Z.H.Z. and M.A.H. supervised the project. D.L.C. and B.-Y.L. prepared protein samples; X.L. made cryoEM grids, performed cryoEM imaging, and data processing; X.L. built the atomic model and illustrated the structures; B.-Y.L. prepared bacterial strains, bacterial two-hybrid assay, and growth competition assays. D.L.C. performed Cu sensitivity studies, protein localization studies, and prepared samples for ICP-MS. X.L., D.L.C., B.-Y.L., M.A.H. and Z.H.Z. interpreted the data and wrote the manuscript; and all authors reviewed and approved the paper.

## Data availability

The cryoEM density map has been deposited in the Electron Microscopy Data Bank under accession codes xxxx. The atomic coordinate has been deposited in the Protein Data Bank under accession codes xxxx.

---

**Supplementary Fig. S1:**
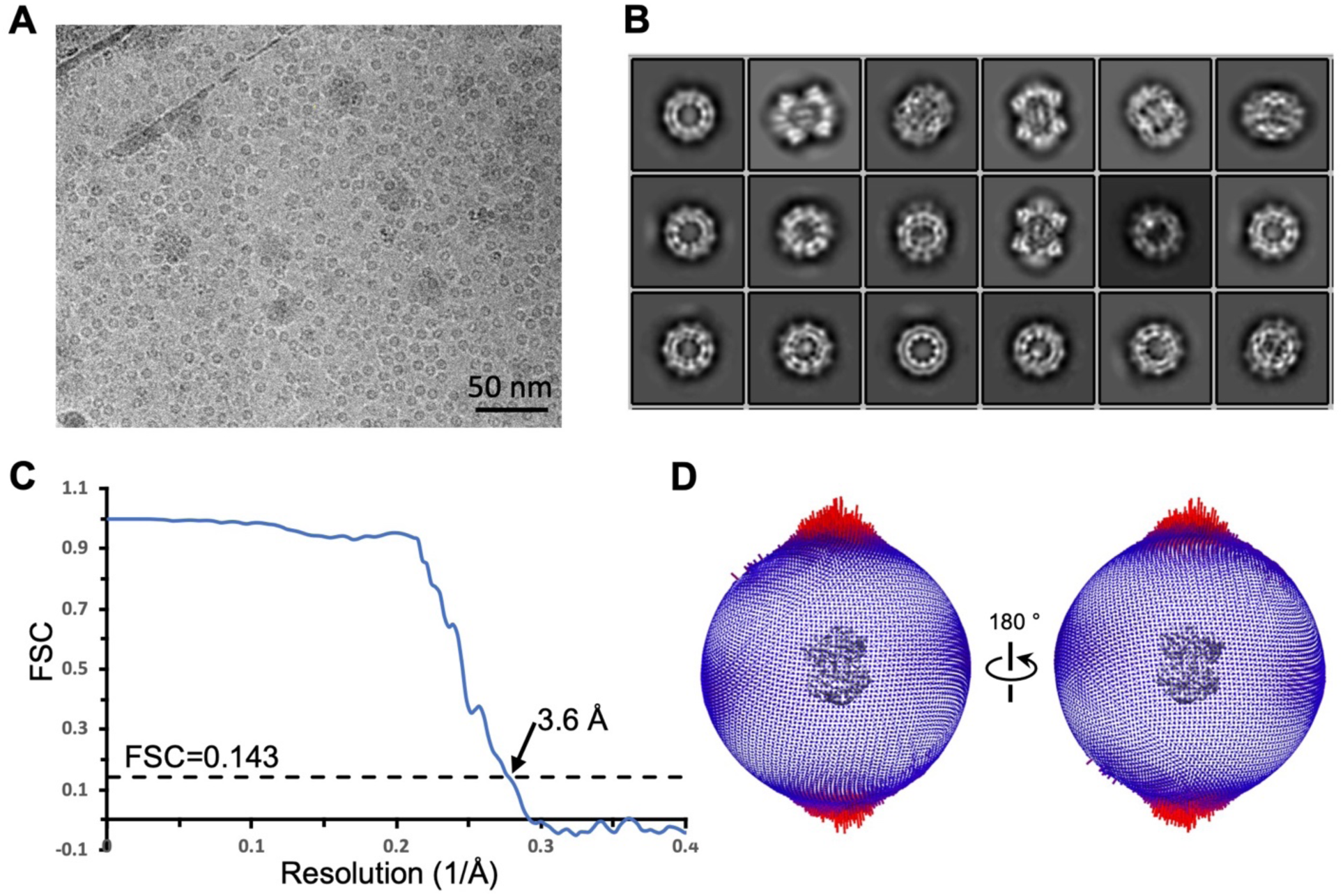
**(A)** Motion-corrected cryoEM micrograph. The scale bar is 50 nm. **(B)** Representative 2D class averages of FPM13 particles. **(C)** Plot of the FSC as a function of the spatial frequency, with resolution indicated. **(D)** Euler angle distributions of FPM13 particles used for the final reconstruction.

**Supplementary Fig. S2:**
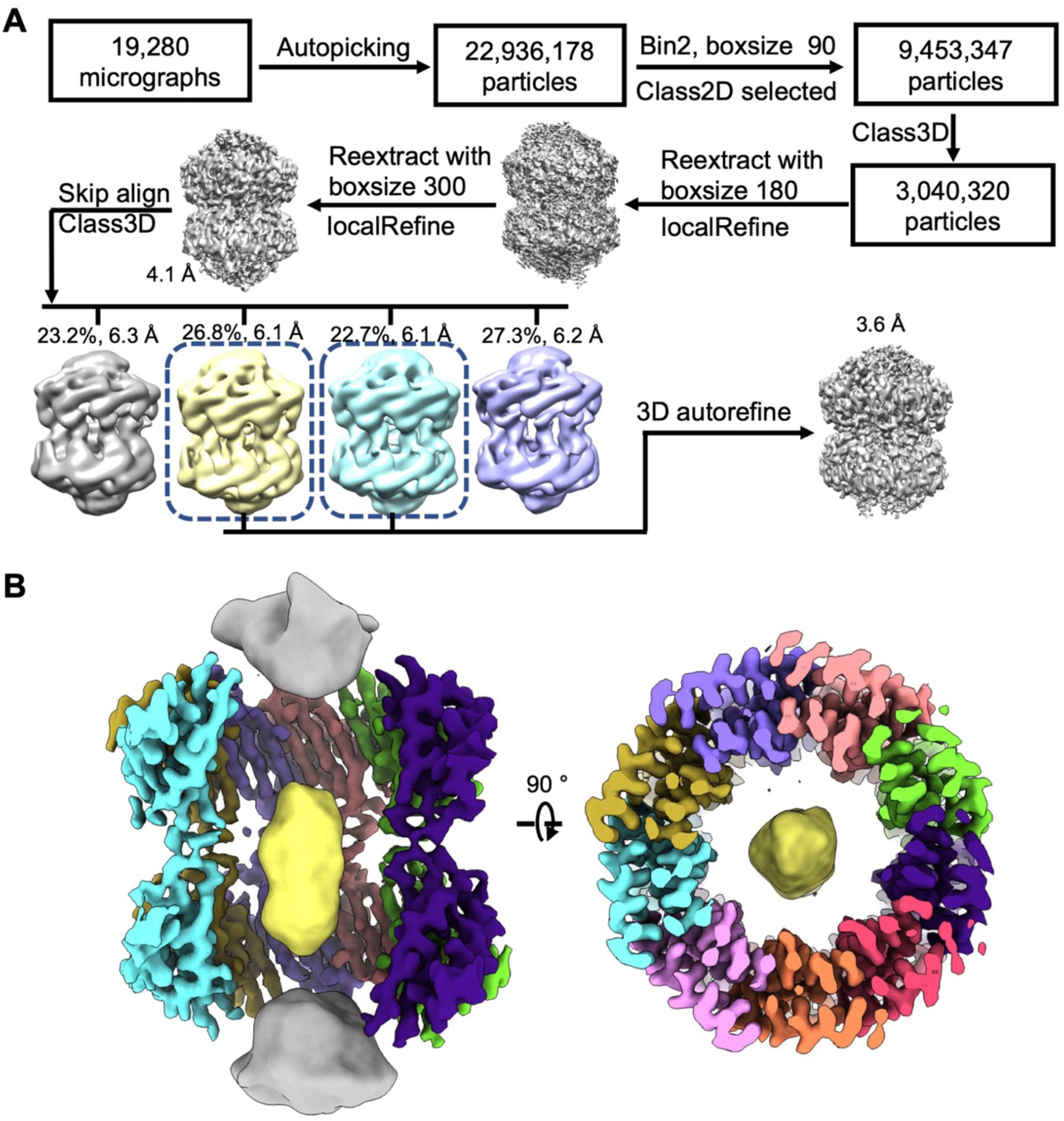
**(A)** CryoEM data processing workflow. Schematic of classification and refinement procedures used to generate the map obtained in this study (see Methods for details). **(B)** Density inside FPM13 shown in yellow.

**Supplementary Fig. S3:**
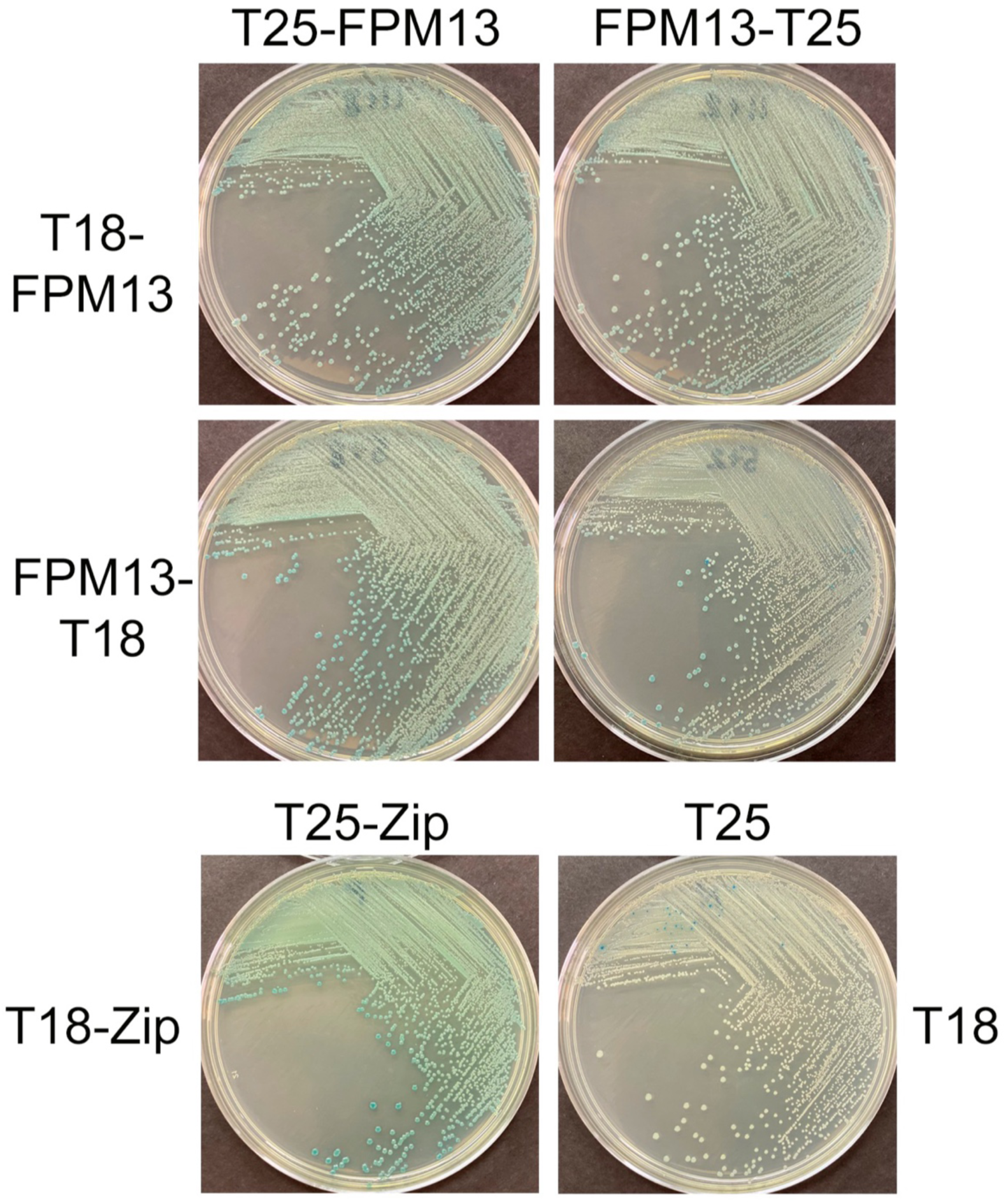
Bacterial 2-hybrid analysis confirms capacity of FPM13 to interact with itself. *E. coli* BTH101 transformed with a pair of compatible plasmids for expression of FPM13 with N- or C-terminal T18 and T25 fusions were streaked onto LB agar plates containing IPTG and X-gal. While a negative interaction causes no change in color, a positive interactions between the T18 and T25 fusion proteins promote cAMP production and beta-galactosidase activity in the *E. coli* and turns the color of the bacterial colonies to blue. *E. coli* transformed with plasmids carrying T18-Zip and T25-Zip fusions served as the positive control. *E. coli* transformed with plasmids carrying T25 and T18 without a fusion partner served as the negative control.

**Supplementary Fig. S4:**
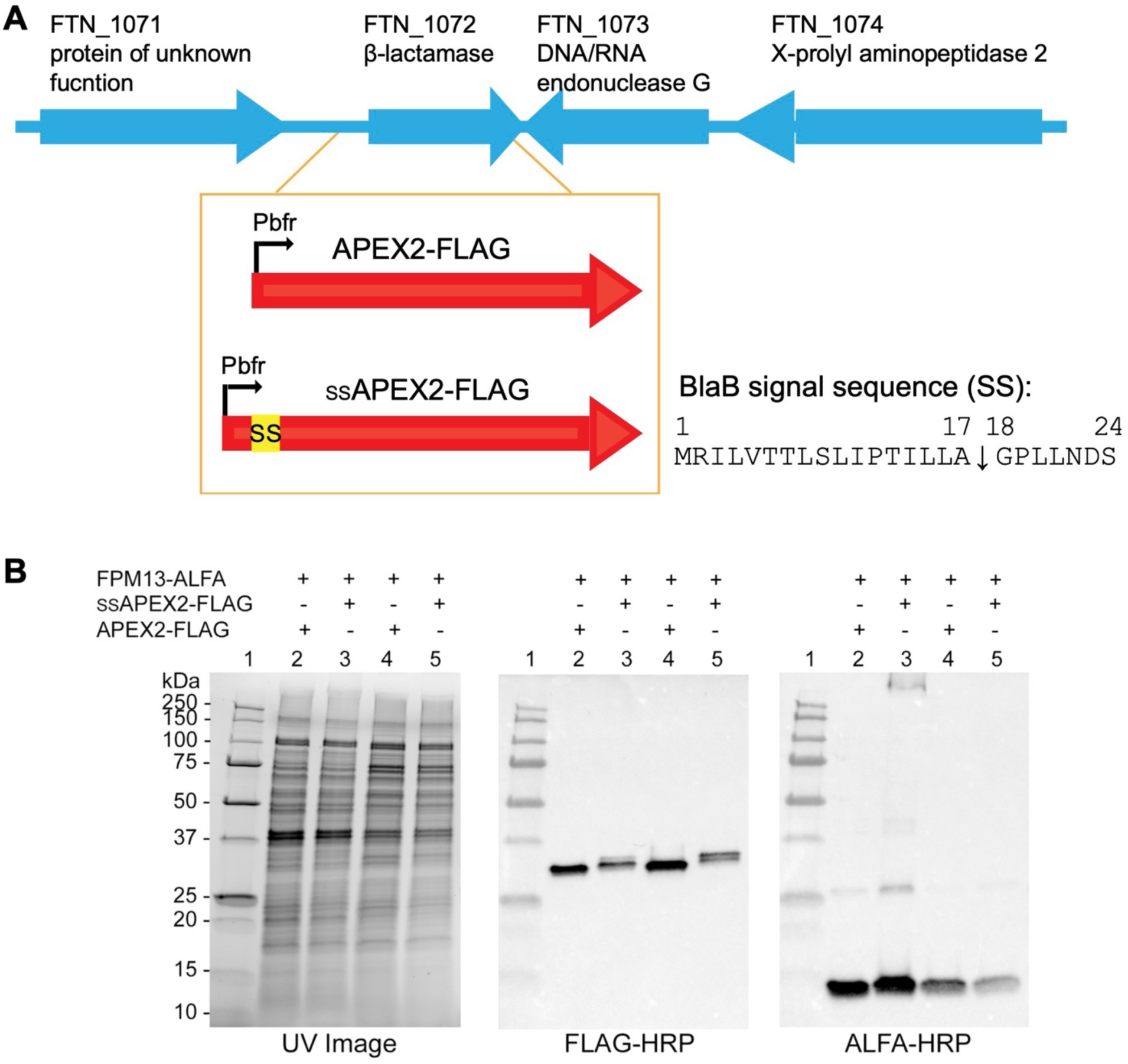
Genomic organization and expression of *F. novicida* strains with ALFA tagged FPM13 and FLAG tagged APEX2 with or without a signal sequence (SS) for secretion. **(A)** Gene cassette of FLAG-tagged APEX2 with or without the first 24 amino acids of the *F. novicida* beta-lactamase (BlaB) driven by the promoter of *F. tularensis* LVS bacterioferritin (bfr) was inserted into the blaB locus. The cleavage site of the BlaB signal peptide sequence is indicated with a downward arrow. **(B)** Proteins from strains grown in TSBC (lanes 2, 3) or TSBC containing 5% KCl (lanes 4, 5) were separated on SDS-PAGE and the protein profiles imaged by stain free UV imaging (left panel). The proteins were transblotted onto nitrocellulose membrane and probed with FLAG-HRP to detect the expression of APEX2 and ssAPEX2 (middle panel) or with ALFA-HRP to detect the expression of FPM13 (right panel). Lane 1, molecular mass standards.

**Supplementary Fig. S5.**
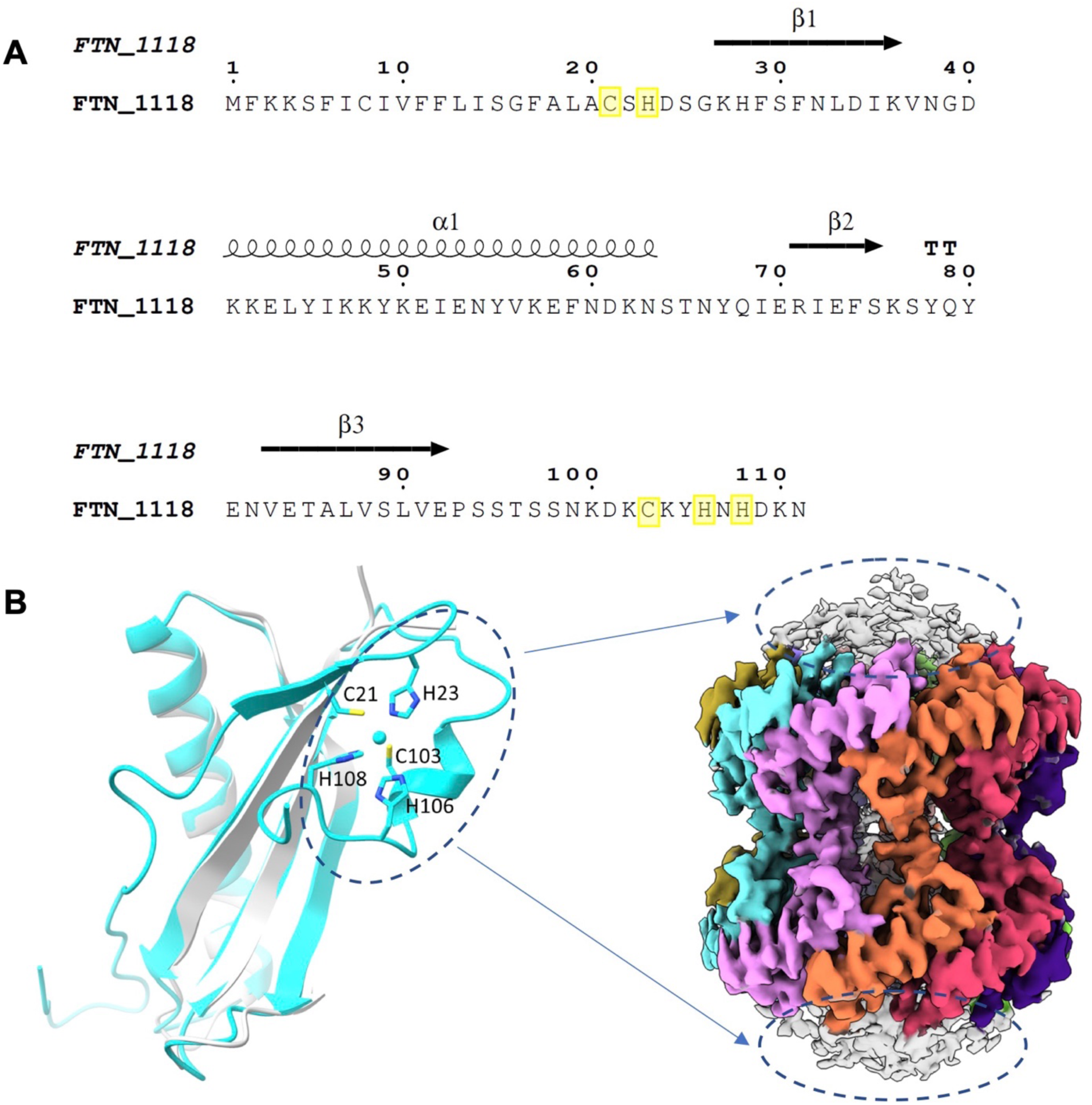
The rationale for FPM13 (FTN_1118) being enriched in the protein purification. **(A)** Sequence and secondary structure of FPM13 protein. **(B)** Chai-1 prediction of FPM13 with Ni^2+^ binding. The ion is shown as a sphere, and key residues are displayed in stick presentation. The binding site is located in the unmodeled top and bottom regions. Key residues involved in ion binding are highlighted in yellow in A.

**Table S1.** CryoEM data collection, refinement and validation statistics.

|  | FPM13<br>EMDB xxxx<br>PDB xxxx |
| --- | --- |
| **Data collection and processing** |  |
| Magnification | 81000 |
| Voltage (kV) | 300 |
| Electron exposure (e<sup>-</sup>/Å<sup>2</sup>) | 50 |
| Defocus range (μm) | -1.8 to -2.6 |
| Pixel size (Å) | 1.1 |
| Symmetry imposed | C1 |
| Particle number | 1503607 |
| Map resolution | 3.6 |
| FSC threshold | 0.143 |
| **Refinement** |  |
| Map sharpening *B* factor (Å<sup>2</sup>) | -202 |
| **Model composition** |  |
| Non-hydrogen atoms | 10602 |
| Protein residues | 1260 |
| Ligand |  |
| ***B* factors (Å<sup>2</sup>)** |  |
| Protein | 56.3 |
| Ligand |  |
| **R.m.s. deviations** |  |
| Bond lengths (Å) | 0.003 |
| Bond angle (°) | 0.645 |
| **Validation** |  |
| MolProbity score | 1.50 |
| Clashscore | 7.36 |
| Poor rotamers (%) | 0.25 |
| **Ramachandran plot** |  |
| Favored (%) | 97.55 |
| Allowed (%) | 2.45 |
| Disallowed (%) | 0 |

**Table S2.**
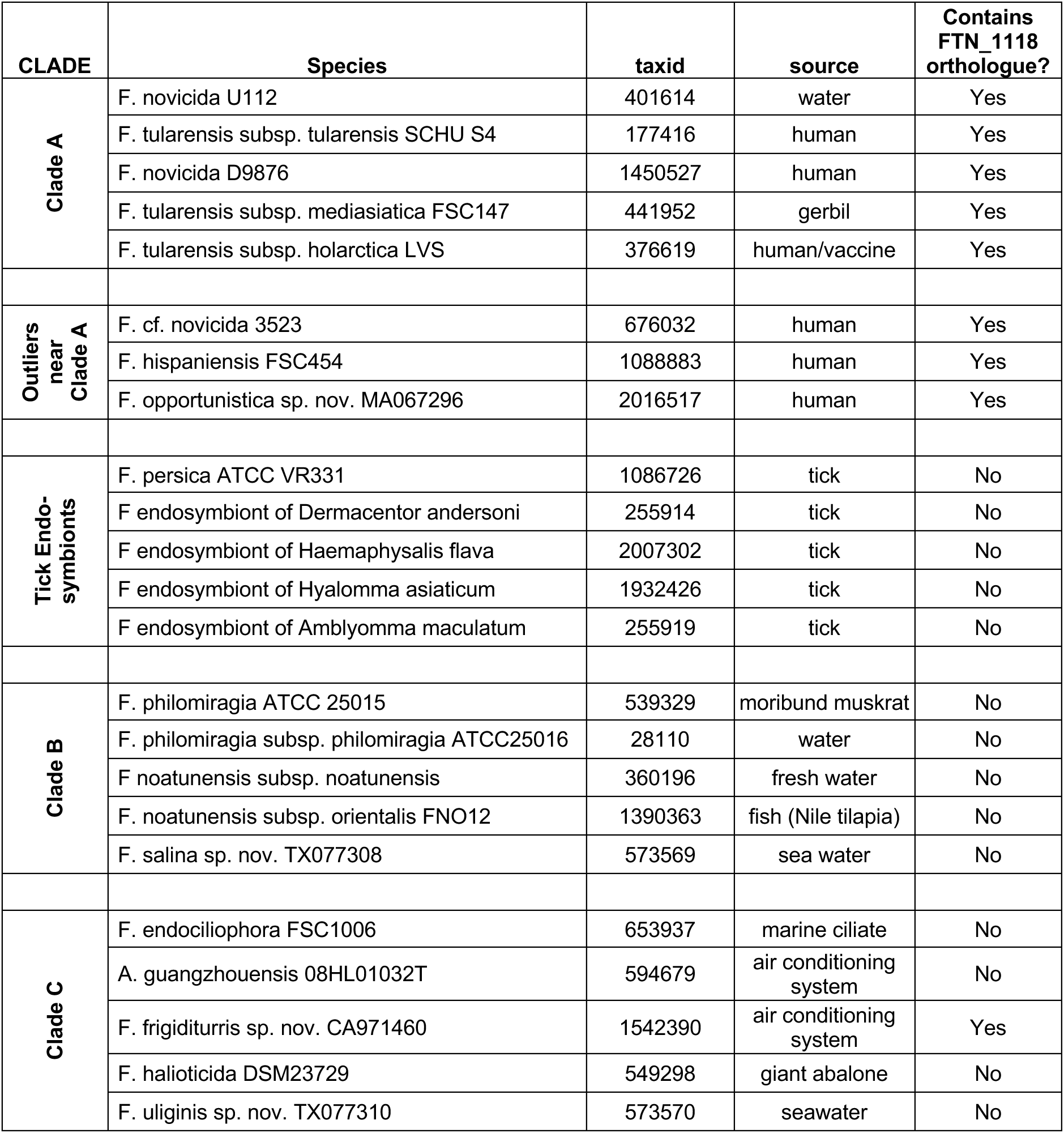
*Francisella* strains classified according to phylogenetic analyses of Duron et al, 2018 and Kuman et al. 2020, indicating presence or absence of FPM13 (FTN_1118) homologs.

| CLADE | Species | taxid | source | Contains<br>FTN_1118<br>orthologue? |
| --- | --- | --- | --- | --- |
| Clade A | F. novicida U112 | 401614 | water | Yes |
|  | F. tularensis subsp. tularensis SCHU S4 | 177416 | human | Yes |
|  | F. novicida D9876 | 1450527 | human | Yes |
|  | F. tularensis subsp. mediasiatica FSC147 | 441952 | gerbil | Yes |
|  | F. tularensis subsp. holarctica LVS | 376619 | human/vaccine | Yes |
| Outliers<br>near<br>Clade A | F. cf. novicida 3523 | 676032 | human | Yes |
|  | F. hispaniensis FSC454 | 1088883 | human | Yes |
|  | F. opportunistica sp. nov. MA067296 | 2016517 | human | Yes |
| Tick Endo-<br>symbionts | F. persica ATCC VR331 | 1086726 | tick | No |
|  | F. endosymbiont of Dermacentor andersoni | 255914 | tick | No |
|  | F. endosymbiont of Haemaphysalis flava | 2007302 | tick | No |
|  | F. endosymbiont of Hyalomma asiaticum | 1932426 | tick | No |
|  | F. endosymbiont of Amblyomma maculatum | 255919 | tick | No |
| Clade B | F. philomiragia ATCC 25015 | 539329 | moribund muskrat | No |
|  | F. philomiragia subsp. philomiragia ATCC25016 | 28110 | water | No |
|  | F. noatunensis subsp. noatunensis | 360196 | fresh water | No |
|  | F. noatunensis subsp. orientalis FNO12 | 1390363 | fish (Nile tilapia) | No |
|  | F. salina sp. nov. TX077308 | 573569 | sea water | No |
| Clade C | F. endociliophora FSC1006 | 653937 | marine ciliate | No |
|  | A. guangzhouensis 08HL01032T | 594679 | air conditioning<br>system | No |
|  | F. frigiditurreis sp. nov. CA971460 | 1542390 | air conditioning<br>system | Yes |
|  | F. haliotica DSM23729 | 549298 | giant abalone | No |
|  | F. uliginis sp. nov. TX077310 | 573570 | seawater | No |

## References

1. Saslaw S, Eigelsbach HT, Wilson HE, Prior JA, Carhart S. Tularemia vaccine study. I. Intracutaneous challenge. Arch Intern Med. 1961;107:689-701. PubMed PMID: 13746668.

2. Saslaw S, Eigelsbach HT, Prior JA, Wilson HE, Carhart S. Tularemia vaccine study. II. Respiratory challenge. Arch Intern Med. 1961;107:702-14. PubMed PMID: 13746667.

3. Sjostedt A. Tularemia: history, epidemiology, pathogen physiology, and clinical manifestations. Ann N Y Acad Sci. 2007;1105:1–29. Epub 20070329. doi: 10.1196/annals.1409.009. PubMed PMID: 17395726.

4. Clemens DL, Lee BY, Horwitz MA. *Francisella tularensis* enters macrophages via a novel process involving pseudopod loops. Infect Immun. 2005;73(9):5892–902. doi: DOI:10.1128/IAI.73.9.5892-5902.2005. PubMed PMID: 16113308; PMCID: 1231130.

5. Clemens DL, Lee BY, Horwitz MA. Virulent and avirulent strains of *Francisella tularensis* prevent acidification and maturation of their phagosomes and escape into the cytoplasm in human macrophages. Infect Immun. 2004;72(6):3204–17. doi: 10.1128/IAI.72.6.3204-3217.2004 72/6/3204 [pii]. PubMed PMID: 15155622; PMCID: 415696.

6. Clemens DL, Horwitz MA. Uptake and intracellular fate of *Francisella tularensis* in human macrophages. Ann N Y Acad Sci. 2007;1105:160–86. doi: doi:10.1196/annals.1409.001. PubMed PMID: 17435118.

7. Barker JR, Chong A, Wehrly TD, Yu JJ, Rodriguez SA, Liu J, Celli J, Arulanandam BP, Klose KE. The Francisella tularensis pathogenicity island encodes a secretion system that is required for phagosome escape and virulence. Mol Microbiol. 2009;74(6):1459–70. Epub 2010/01/08. doi: 10.1111/j.1365-2958.2009.06947.x. PubMed PMID: 20054881; PMCID: PMC2814410.

8. Clemens DL, Lee BY, Horwitz MA. The *Francisella* Type VI Secretion System. Front Cell Infect Microbiol. 2018;8:121. doi: 10.3389/fcimb.2018.00121. PubMed PMID: 29740542; PMCID: 5924787.

9. Larson CL, Wicht W, Jellison WL. A new organism resembling P. tularensis isolated from water. Public Health Rep (1896). 1955;70(3):253–8. PubMed PMID: 14357545; PMCID: PMC2024510.

10. Brett ME, Respicio-Kingry LB, Yendell S, Ratard R, Hand J, Balsamo G, Scott-Waldron C, O’Neal C, Kidwell D, Yockey B, Singh P, Carpenter J, Hill V, Petersen JM, Mead P. Outbreak of Francisella novicida Bacteremia Among Inmates at a Louisiana Correctional Facility. Clin Infect Dis. 2014;59(6):826–33. doi: 10.1093/cid/ciu430.

11. Pfab J, Phan NM, Si D. DeepTracer for fast de novo cryo-EM protein structure modeling and special studies on CoV-related complexes. Proc Natl Acad Sci U S A. 2021;118(2). doi: 10.1073/pnas.2017525118. PubMed PMID: 33361332; PMCID: PMC7812826.

12. Ho CM, Li X, Lai M, Terwilliger TC, Beck JR, Wohlschlegel J, Goldberg DE, Fitzpatrick AWP, Zhou ZH. Bottom-up structural proteomics: cryoEM of protein complexes enriched from the cellular milieu. Nat Methods. 2020;17(1):79–85. Epub 20191125. doi: 10.1038/s41592-019-0637-y. PubMed PMID: 31768063; PMCID: PMC7494424.

13. Holm L, Rosenstrom P. Dali server: conservation mapping in 3D. Nucleic Acids Res. 2010;38(Web Server issue):W545–9. doi: 10.1093/nar/gkq366. PubMed PMID: 20457744; PMCID: 2896194.

14. Yu NY, Wagner JR, Laird MR, Melli G, Rey S, Lo R, Dao P, Sahinalp SC, Ester M, Foster LJ, Brinkman FS. PSORTb 3.0: improved protein subcellular localization prediction with refined localization subcategories and predictive capabilities for all prokaryotes. Bioinformatics. 2010;26(13):1608–15. Epub 20100513. doi: 10.1093/bioinformatics/btq249. PubMed PMID: 20472543; PMCID: PMC2887053.

15. Consortium TU. UniProt: the Universal Protein Knowledgebase in 2025. Nucleic Acids Res. 2024;53(D1):D609–D17. doi: 10.1093/nar/gkae1010.

16. Ganapathy US, Bai L, Wei L, Eckartt KA, Lett CM, Previti ML, Carrico IS, Seeliger JC. Compartment-Specific Labeling of Bacterial Periplasmic Proteins by Peroxidase-Mediated Biotinylation. ACS infectious diseases. 2018;4(6):918–25. Epub 20180507. doi: 10.1021/acsinfecdis.8b00044. PubMed PMID: 29708735; PMCID: PMC6767932.

17. Hwang J, Espenshade PJ. Proximity-dependent biotin labelling in yeast using the engineered ascorbate peroxidase APEX2. Biochem J. 2016;473(16):2463–9. Epub 20160607. doi: 10.1042/bcj20160106. PubMed PMID: 27274088; PMCID: PMC5290329.

18. Brusca JS, Radolf JD. Isolation of integral membrane proteins by phase partitioning with Triton X-114. Methods Enzymol. 1994;228:182–93. doi: 10.1016/0076-6879(94)28019-3. PubMed PMID: 8047007.

19. Robertson GT, Child R, Ingle C, Celli J, Norgard MV. IglE is an outer membrane-associated lipoprotein essential for intracellular survival and murine virulence of type A *Francisella tularensis*. Infect Immun. 2013;81(11):4026–40. Epub 2013/08/21. doi: 10.1128/IAI.00595-13 IAI.00595-13 [pii]. PubMed PMID: 23959721; PMCID: 3811846.

20. Nguyen JQ, Gilley RP, Zogaj X, Rodriguez SA, Klose KE. Lipidation of the FPI protein IglE contributes to *Francisella tularensis* ssp. *novicida* intramacrophage replication and virulence. Pathogens and disease. 2014;72(1):10–8. doi: 10.1111/2049-632X.12167. PubMed PMID: 24616435; PMCID: 4160424.

21. Duron O, Morel O, Noel V, Buysse M, Binetruy F, Lancelot R, Loire E, Menard C, Bouchez O, Vavre F, Vial L. Tick-Bacteria Mutualism Depends on B Vitamin Synthesis Pathways. Curr Biol. 2018;28(12):1896–902 e5. Epub 20180531. doi: 10.1016/j.cub.2018.04.038. PubMed PMID: 29861133.

22. Kumar R, Broms JE, Sjostedt A. Exploring the Diversity Within the Genus Francisella - An Integrated Pan-Genome and Genome-Mining Approach. Front Microbiol. 2020;11:1928. Epub 20200811. doi: 10.3389/fmicb.2020.01928. PubMed PMID: 32849479; PMCID: PMC7431613.

23. Camacho C, Coulouris G, Avagyan V, Ma N, Papadopoulos J, Bealer K, Madden TL. BLAST+: architecture and applications. BMC Bioinformatics. 2009;10:421. Epub 20091215. doi: 10.1186/1471-2105-10-421. PubMed PMID: 20003500; PMCID: PMC2803857.

24. Djoko KY, Ong CL, Walker MJ, McEwan AG. The Role of Copper and Zinc Toxicity in Innate Immune Defense against Bacterial Pathogens. J Biol Chem. 2015;290(31):18954–61. Epub 20150608. doi: 10.1074/jbc.R115.647099. PubMed PMID: 26055706; PMCID: PMC4521016.

25. German N, Doyscher D, Rensing C. Bacterial killing in macrophages and amoeba: do they all use a brass dagger? Future microbiology. 2013;8(10):1257–64. doi: 10.2217/fmb.13.100. PubMed PMID: 24059917.

26. LoVullo ED, Sherrill LA, Perez LL, Pavelka MS, Jr. Genetic tools for highly pathogenic Francisella tularensis subsp. tularensis. Microbiology. 2006;152(Pt 11):3425–35. doi: 152/11/3425 [pii] 10.1099/mic.0.29121-0. PubMed PMID: 17074911.

27. Lam SS, Martell JD, Kamer KJ, Deerinck TJ, Ellisman MH, Mootha VK, Ting AY. Directed evolution of APEX2 for electron microscopy and proximity labeling. Nat Methods. 2015;12(1):51–4. Epub 20141124. doi: 10.1038/nmeth.3179. PubMed PMID: 25419960; PMCID: PMC4296904.

28. Liu X, Clemens DL, Lee BY, Aguirre R, Horwitz MA, Zhou ZH. Structure, identification and characterization of the RibD-enolase complex in Francisella. bioRxiv. 2025. Epub 20250303. doi: 10.1101/2025.03.02.641097. PubMed PMID: 40093042; PMCID: PMC11908141.

29. Mastronarde DN. Automated electron microscope tomography using robust prediction of specimen movements. J Struct Biol. 2005;152(1):36–51. doi: 10.1016/j.jsb.2005.07.007. PubMed PMID: 16182563.

30. Zheng SQ, Palovcak E, Armache JP, Verba KA, Cheng Y, Agard DA. MotionCor2: anisotropic correction of beam-induced motion for improved cryo-electron microscopy. Nat Methods. 2017;14(4):331–2. Epub 20170227. doi: 10.1038/nmeth.4193. PubMed PMID: 28250466; PMCID: PMC5494038.

31. Rohou A, Grigorieff N. CTFFIND4: Fast and accurate defocus estimation from electron micrographs. Journal of structural biology. 2015;192(2):216–21. doi: 10.1016/j.jsb.2015.08.008. PubMed PMID: 26278980.

32. Scheres SH. Processing of Structurally Heterogeneous Cryo-EM Data in RELION. Methods Enzymol. 2016;579:125–57. Epub 2016/08/31. doi: 10.1016/bs.mie.2016.04.012. PubMed PMID: 27572726.

33. Scheres SH. RELION: implementation of a Bayesian approach to cryo-EM structure determination. J Struct Biol. 2012;180(3):519–30. Epub 2012/09/25. doi: 10.1016/j.jsb.2012.09.006. PubMed PMID: 23000701; PMCID: PMC3690530.

34. Emsley P, Cowtan K. Coot: model-building tools for molecular graphics. Acta Crystallogr D Biol Crystallogr. 2004;60(Pt 12 Pt 1):2126–32. Epub 2004/12/02. doi: 10.1107/S0907444904019158. PubMed PMID: 15572765.

35. Adams PD, Afonine PV, Bunkoczi G, Chen VB, Davis IW, Echols N, Headd JJ, Hung LW, Kapral GJ, Grosse-Kunstleve RW, McCoy AJ, Moriarty NW, Oeffner R, Read RJ, Richardson DC, Richardson JS, Terwilliger TC, Zwart PH. PHENIX: a comprehensive Python-based system for macromolecular structure solution. Acta crystallographica Section D, Biological crystallography. 2010;66(Pt 2):213–21. doi: 10.1107/S0907444909052925. PubMed PMID: 20124702; PMCID: 2815670.

36. Pettersen EF, Goddard TD, Huang CC, Couch GS, Greenblatt DM, Meng EC, Ferrin TE. UCSF Chimera--a visualization system for exploratory research and analysis. J Comput Chem. 2004;25(13):1605–12. Epub 2004/07/21. doi: 10.1002/jcc.20084. PubMed PMID: 15264254.

37. Goddard TD, Huang CC, Meng EC, Pettersen EF, Couch GS, Morris JH, Ferrin TE. UCSF ChimeraX: Meeting modern challenges in visualization and analysis. Protein Sci. 2018;27(1):14–25. Epub 2017/07/16. doi: 10.1002/pro.3235. PubMed PMID: 28710774; PMCID: PMC5734306.

38. Lee BY, Horwitz MA, Clemens DL. Identification, recombinant expression, immunolocalization in macrophages, and T-cell responsiveness of the major extracellular proteins of *Francisella tularensis*. Infect Immun. 2006;74(7):4002–13. doi: 74/7/4002 [pii] 10.1128/IAI.00257-06. PubMed PMID: 16790773.

39. Chamberlain RE. Evaluation of Live Tularemia Vaccine Prepared in a Chemically Defined Medium. Appl Microbiol. 1965;13:232–5. Epub 1965/03/01. PubMed PMID: 14325885; PMCID: 1058227.

